# Desynchrony between events triggers a compensatory delay during *C. elegans* development

**DOI:** 10.1101/2024.03.31.587169

**Authors:** Francisco J. Romero-Expósito, Almudena Moreno-Rivero, Marta Muñoz-Barrera, Nicola Gritti, Francesca Sartor, Martha Merrow, Jeroen S. van Zon, Alejandro Mata-Cabana, María Olmedo

## Abstract

In multicellular organisms, development entails the progression of diverse cellular processes that need to be temporally coordinated. During *C. elegans* postembryonic development, the events of molting and cell divisions progress in parallel during four larval stages and are modulated by external cues such as temperature and nutrient availability. While seam cell divisions occur predominantly before ecdysis of the cuticle, the order of the events can change, suggesting that they are controlled by independent mechanisms. Here, we have analyzed the impact of reduced insulin signaling in molting and in stage-specific cell divisions. We find that reduced insulin signaling in the *daf-2(e1370)* allele delays both of these events but has a larger impact on the timing of cell divisions, thus increasing the desynchrony between the events. The relative delay in seam cell divisions leads to a delay in the initiation of the subsequent stage of the molting program, providing a mechanism for resynchronization of developmental processes to the beginning of the larval stages.

## INTRODUCTION

Development is a temporal program made up of different modules that typically proceed in a precise series of events. To achieve reproducible development, organisms must integrate information from intrinsic developmental timers with external cues, such as changes in nutrition and temperature. During the postembryonic development of *C. elegans*, each larval stage is defined by distinct cellular events of division, migration, and differentiation. These events have been meticulously traced due to the highly consistent nature of the process. Seam cells divide asymmetrically, approximately at the beginning of each larval stage. At the beginning of the second larval stage (L2) there is, in addition, a symmetric division before the asymmetric [1]. The heterochronic pathway specifies what cell division events take place during each stage (reviewed in [2]). Additionally, across the four larval stages (L1–L4), development proceeds through a repeated cycle in which each stage includes an intermolt period (I1–I4) and a subsequent molt (M1–M4). Molting is a period defined by the generation of a new cuticle and ends with the shedding of the old cuticle, a process known as ecdysis [3]. Oscillatory gene expression of > 3,700 genes is coupled to the process of molting [4]. Cell divisions and molting show an apparent coordination through postembryonic development, with events taking place in an orderly manner. However, the timing of the events of cell division and molting seem to be controlled by independent timers that can be uncoupled [5], leading to alterations in the characteristic sequence [6].

Diverse environmental factors, including temperature and nutrition, impact the speed of postembryonic development in a stage-dependent manner [6–9]. One of the major pathways controlling of *C. elegans* development is that for Insulin/IGF-1 signaling (IIS). This nutrient sensing pathway controls lifespan and developmental progression and is conserved across metazoans. In *C. elegans*, the gene *daf-2* encodes the sole insulin receptor DAF-2/IR. Signaling through the pathway occurs to a large extent through the phosphatidylinositol-3-OH kinase (PI3K) AGE-1, and the serine/threonine kinases PDK-1, AKT-1, and AKT-2. These proteins regulate the activity of the transcription factor (TF) DAF-16/FOXO. Reduced DAF-2 function leads to increased lifespan in a DAF-16 dependent manner, suggesting that this TF is the main effector of the pathway (reviewed in [10]). Hypomorphic *daf-2* alleles also show stage-specific developmental delays [7,9], but the impact of reduced insulin signaling in the timing of postembryonic cell divisions had not been tested before.

We have applied a time-resolved analysis of the molting process to analyze mutants of the insulin signaling pathway, unveiling a novel feature of development. The hypomorphic *daf-2(e1370)* mutant shows long lags in the initiation of intermolts, especially after finishing the first molt (named L2lag as it delays the L2 intermolt). This *daf-2(e1370)* phenotype is partially dependent on *daf-16* and *daf-18*. We then analyzed seam cell division events and observed that *daf-2(e1370)* also delays the timing of seam cell divisions. However, the delay in cell divisions is greater than that of molting, increasing desynchrony between ecdysis and cell divisions at the end of L1. By combining two reporter systems to monitor molting and seam cell division in the same animal, we observed that larvae in L2lag present a delay in seam cell division relative to ecdysis. This observation links L2lag to the desynchrony between developmental events. To show causality, we experimentally changed the timing of cell divisions relative to ecdysis and tested the effect on L2lag. Our results suggest a model in which delayed cell divisions can be compensated by delaying the initiation of the following stage, so that desynchrony does not amplify over the process of development.

## RESULTS

### A lag phase in the initiation of larval stages

Analysis of postembryonic development in *C. elegans* has often relied on observation of morphological changes. Here, we applied a high-throughput, quantitative method to measure development [11] that allows unveiling unknown complexities on the timing of the different larval stages [7,12,13]. This method relies on the use of the enzyme Luciferase, which is constitutively expressed in the *C. elegans* reporter strain *sevIs1 [Psur-5::luc+::gfp]* [11]. Luciferase catalyzes the oxidation of Luciferin, in a reaction that emits light [14]. We provide Luciferin in the media of the developing larvae, so that luciferin intake results in emission of light. At the beginning of the molts, larvae produce a cuticular plug that blocks the mouth, preventing the entry of luciferin. As a consequence, the emission of light decreases rapidly, generating a characteristic profile that is used to calculate the duration of each larval stage, allowing identification of intermolts (I) and molts (M) (Fig. 1A) [11].

**Figure 1.**
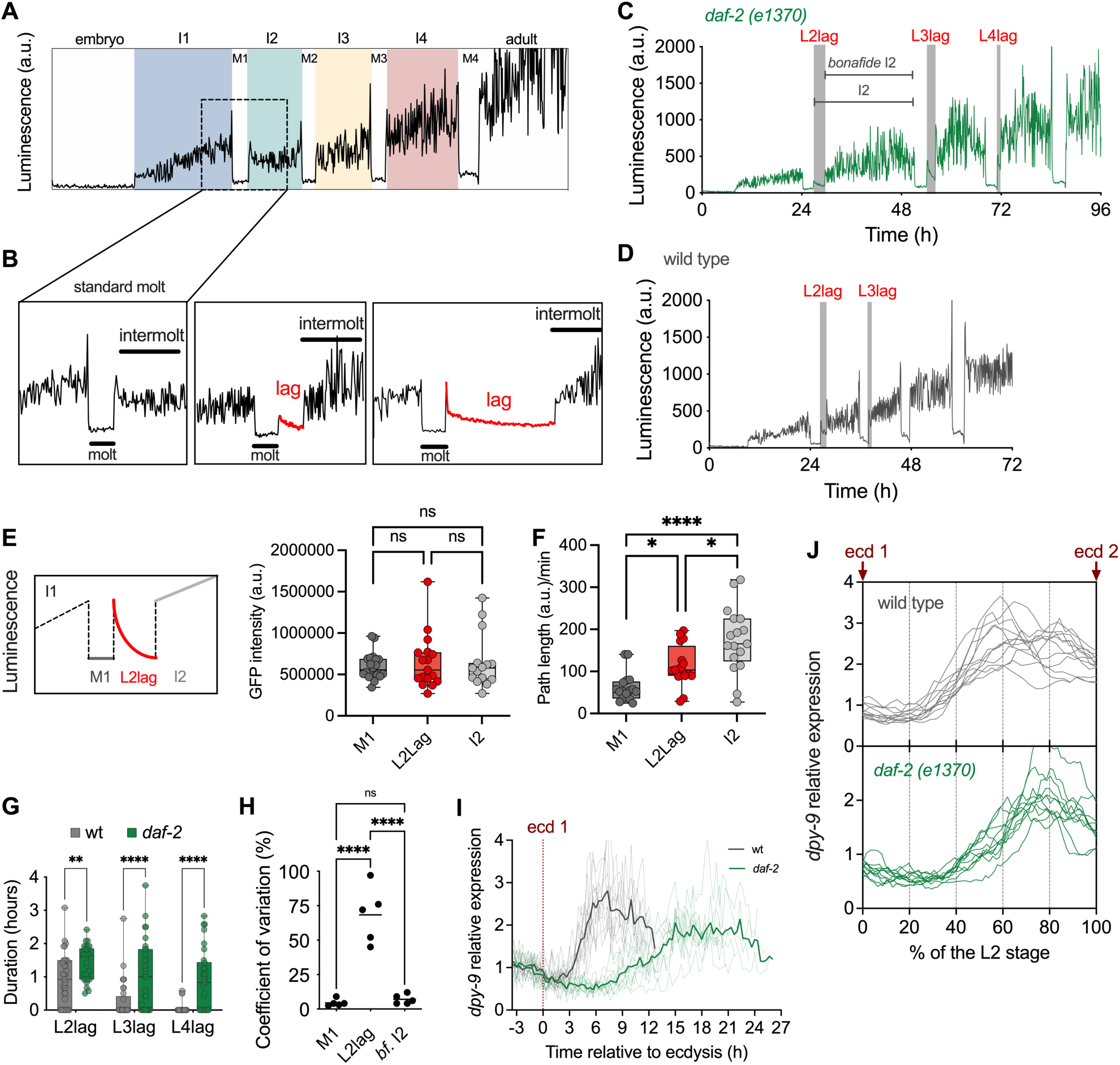
*daf-2(e1370)* shows a novel developmental phenotype. **(A)** Representative profile of luminescence over postembryonic development **(B)** Zoom in of a standard molt (left) and profiles showing representative short and long lags (center and right). **(C)** Representative profile of luminescence of a *daf-2(e1370)* larvae, showing L2lag, L3lag and L4lag. **(D)** Representative profile of luminescence of a single wild-type larvae, indicating the presence of L2lag. **(E)** Diagram showing the moment of collection and corresponding values of GFP intensity (a.u.). **(F)** Path length per minute (a.u.) for the same categories of larvae as in E. **(G)** Duration of the lag in the wild type and *daf-2(e1370)* larvae. Statistical differences show Fisheŕs LSD between strains after two-way ANOVA, ** p<0.01, **** p<0.0001. The dots show individual larvae in 3 independent experiments. **(H)** Coefficient of variation per experiment for the duration of M1, L2lag and *bonafide* I2 for 56 *daf-2* mutant larvae in 5 independent experiments. Statistical analysis shows one-way ANOVA followed by Tukeýs multiple comparison **** p<0.0001. **(I)** Expression of the molting reporter *dpy-9p::*GFP around ecdysis 1 for individual larvae of the wild type and *daf-2(e1370)* the mutant. Thin traces represent the values of individual larvae while thick traces represent the averages. The signal is normalized to the average signal before ecdysis. **(J)** Expression of the same individual larva after rescaling to the total duration of the L2 stage for each individual larva. A complete report of replicates and statistics results is shown in S2 Table and S1 File.

In the analysis of developmental phenotypes of the *daf-2(e1370)* allele, we observed a novel feature of postembryonic development. In addition to the slow development previously described for this mutant, we detected that, after the end of the molt, many larvae present a lag phase before the resumption of the sustained high signal that is a feature of the intermolts (Fig. 1B, middle and right). To quantify this feature of development, we defined the periods L2lag, L3lag, and L4lag, which respectively appeared at the beginning of the L2, L3, and L4 (Fig. 1C). While initially most evident in *daf-2* mutants, detailed analysis of the wild-type luminescence traces also allowed to identify Lag periods (Fig. 1D). To analyze whether the reduction of the luminescence signal during the lags stems from a reduced expression of the luciferase protein under the promoter of *sur-5*, we measured GFP in animals collected during the first molt (M1), L2lag, and the second intermolt (I2). The three categories showed similar levels of GFP expression, suggesting that the decrease of the signal is not a consequence of reduced luciferase expression (Fig. 1E). In these experiments we could visualize larvae in L2lag, that do not show any feature that distinguished them from the other categories. Since the level of luminescence signal during L2lag is similar to that of molts, and molts involve behavioral quiescence, we tested this variable in the same categories as before. Larvae in L2lag move significantly more than those in M1, and less than larva in the second intermolt (Fig. 1F). Throughout these experiments, we could visualize larvae in the L2lag stage. These larvae showed no apparent morphological differences relative to the other stage and did not present any cuticular remnants. Furthermore, to confirm that L2lag is not construct specific, we analyzed another luciferase strain with a single-copy insertion and a different promoted driving the expression of luciferase [4]. L2lag and larval stages show the same duration in both the wild-type and *daf-2(e1370)* backgrounds (Fig. S1A).

We then compared the levels of lags in the wild type and the *daf-2(e1370)* mutant. The absence of this feature in some individual larvae is denoted by a duration of “0” hours. The lags in the wild-type strain appeared at a lower frequency (approximately in 60% of wild type larvae vs. 90% of *daf-2*) and with shorter duration than in *daf-2* (Fig. 1G). Here, we focus our analysis on L2lag, as this is the stage with the longest average duration, especially for the wild-type strain. We compared the L2lag with the previous and subsequent stages, M1 and *bonafide* I2 respectively, with *bonafide* I2 being the duration of the second intermolt starting after the L2lag period (Fig. 1C). A key feature of L2lag is that, unlike molts and intermolts, the duration of these periods shows high inter-individual variability. The coefficient of variation increases around tenfold for L2lag compared to that of the previous and following stages (Fig. 1 H). We also calculated, for each larva and for the different larval stages, the duration relative to the average (in fold change). This value is close to 1 for M1 and *bonafide* I2, reflecting the low variability in the duration of these stages, but it ranges from 0 to 4 for the L2lag stage (Fig. S1B).

When we compare the relative duration of L2lag and *bf*. I2 in individual larvae, we observed no correlation between these values (Fig. S1C). We also checked the correlation between the duration of each of the stages of development for the wild type and *daf-2*. As previously shown by us and others, larvae show some degree of correlation in the duration of most stages [7,15], but the duration of the lag periods (L2lag, L3lag and L4lag) showed, in general, no correlation with the duration of any of the stages (Fig. S1 D-E). These results show that the duration of the lags does not reflect the interindividual changes in the duration of other developmental stages, that is, L2lag is not a consequence of the presence of larvae with overall faster or slower development. Indeed, for the larvae that present L2lag, if we represent the duration of the complete intermolt, strictly considering the time between molts, its duration is significantly higher than that of larvae that do not enter this stage. By subtracting the duration of the L2lags, we obtain a *bonafide* I2 intermolt duration that is statistically not different from the average I2 (Fig. S1F). All the previous results support that L2lag is a period that is added to the (*bonafide)* intermolt, as opposed to a feature of the luminometry signal accommodated during the duration of the regular intermolt.

Lag occurs at the beginning of larval stages, the same moment when a nutritional checkpoint was previously observed [16], and also coincides with the phase of arrest of developmental oscillations in gene expression [4]. We therefore tested the expression of the gene *dpy-9* as a reporter of these oscillations [4]. We performed time-lapse microscopy of single larvae contained in 275×275×30 μm hydrogel microchambers [17] to monitor the expression of *dpy-9* in the wild type and *daf-2* mutants during the complete L2 stage. On average, the increase of *dpy-9* signal was delayed in the *daf-2* mutant (Fig. 1I). However, as the duration of the L2 stage is different between the wild type and the *daf-2* mutant, and among individual larvae of the same genotype, we plotted the data rescaled to the duration of each individual larva. Even after rescaling, we found that the signal increased later in *daf-2* mutants (Fig. 1J). This data supports the existence of delay in the expression of oscillatory gene expression at the beginning of L2 in *daf-2* mutants. All results together show that the L2lag has features that are different to that of molts and intermolts, with a higher CV and a lack of correlation with the duration of other developmental stages. Furthermore, this lag could coincide with a delay in the initiation of the gene expression characteristic of the second larval stage.

### The L2lag phenotype of *daf-2(e1370)* is partially independent from the rest of IIS

To investigate the regulation of L2lag, we focused on two relevant regulators of IIS. Most *daf-2* phenotypes are mediated by the activation of DAF-16 (Fig. 2A), and they are rescued by the null mutation *daf-16(mu86)*. Thus, phenotypes observed in *daf-2* mutants as constitutive dauer formation, lifespan extension, and the increased duration of the L2 stage, are absent in the double mutant *daf-2(e1370)*;*daf-16(mu86)* [9,18,19]. As expected, we observed that both *daf-16* and the double *daf-2;daf-16* mutants show a duration of the second *bonafide* intermolt similar to the wild-type strain (Fig. 2B). Therefore, the extension of this stage in the *daf-2(e1370)* mutant requires the function of DAF-16. When we quantified L2lag, we observed that *daf-16* has similar levels to the wild-type strain but the double mutant *daf-2;daf-16* shows L2lag levels that are intermediate to the *daf-2* and the *daf-16* mutant (Fig. 2C). The *daf-16* mutation also reduced the levels of L3lag and L4 lag of *daf-2* (Fig. S2A,B). These results show that part of the impact of the *daf-2* mutation is independent of DAF-16. Also, it reinforces the idea that the L2lag phenotype is independent of other developmental phenotypes of *daf-2(e1370)*.

**Figure 2.**
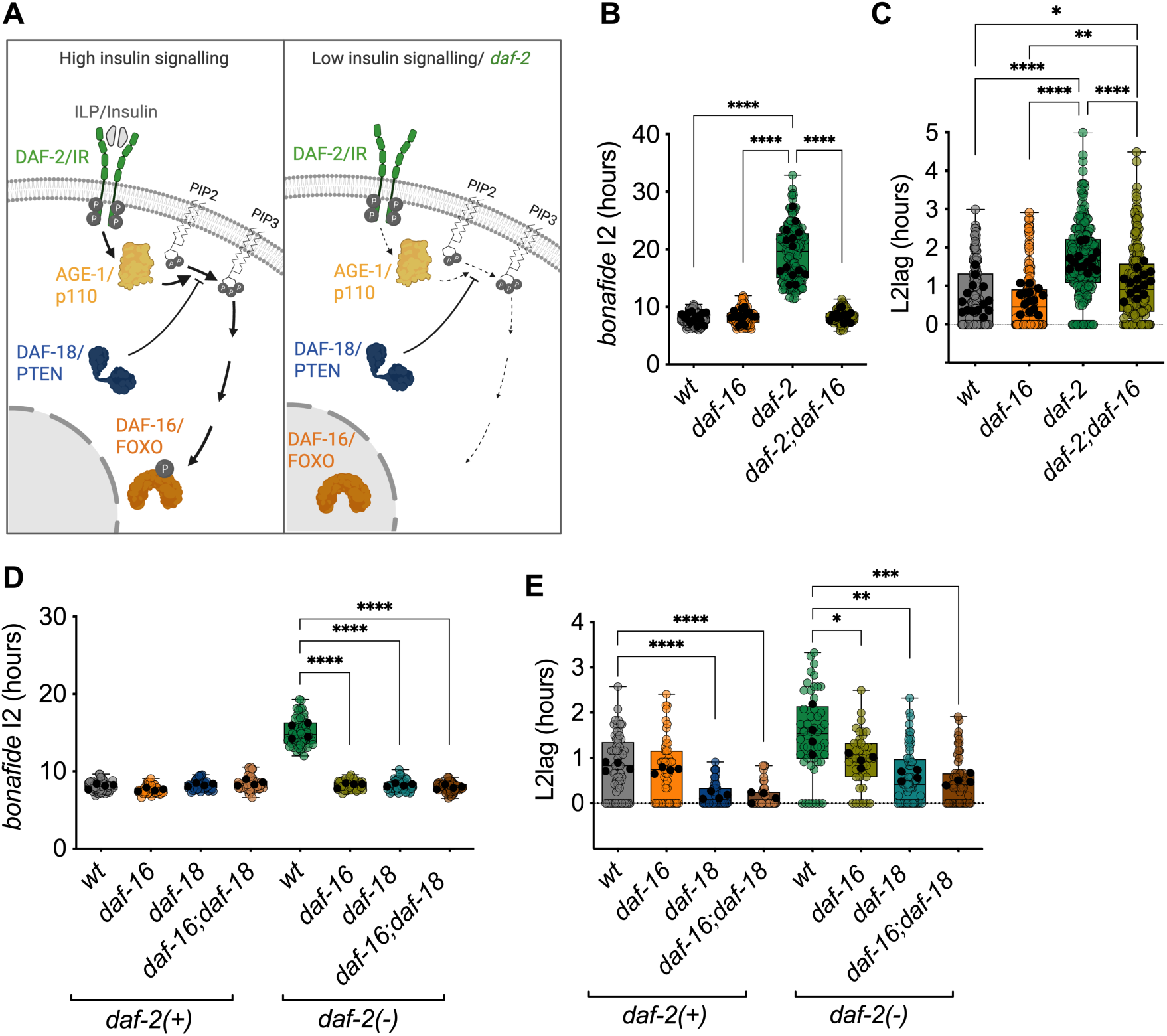
*daf-2(e1370)* leads to L2lag by DAF-16 and DAF-18 dependent and independent pathways. **(A)** Representation of PI3K-dependent IIS in conditions of high insulin signaling. Under low insulin signaling, as in a *daf-2* mutant, signaling through the pathway is reduced, maintaining DAF-16 in a non-phosphorylated state that results in translocation to the nucleus. **(B, C)** Duration of the *bonafide* I2 (B), and L2lag (C), in the wild-type strain, *daf-16(mu86)*, *daf-2(e1370)*, and the double mutant *daf-2;daf-16*, from 8 independent experiments. **(D, E)** Duration of the *bonafide* I2 (D), and L2lag (E), for the wild type, *daf-2*, *daf-16*, *daf-18*, and the respective double and triple mutants. Colored circles show values for individual larvae and black dots show the average for each experiment. For all panels, we performed One-way ANOVA followed by Tukeýs multiple comparisons to test differences between all strains. For B and C all significant differences are shown. For D and E all significant differences within each of the *daf-2* backgrounds are shown. In all cases, * p<0.05 ** p<0.01, ***p<0.001, **** p<0.0001. Complete report of replicates and statistics results is shown in S2 Table and S1 File.

Looking for additional regulators of the process, we focused on DAF-18, the *C. elegans* homologue of the tumor suppressor PTEN, which counteracts insulin signaling by dephosphorylating PIP_3_, opposing the effect of AGE-1 [20] (Fig. 2A). Furthermore, PTEN can dephosphorylate the Insulin Receptor (IR) and the IR inhibits PTEN [21]. We tested the effect of the deletion allele *daf-18(ok480)* in the developmental phenotypes observed previously. The *daf-18* mutation does not provoke any changes in the duration of the second intermolt in a background wildtype for *daf-2* [*daf-2 (+)*] and completely suppresses the *daf-2* mutant phenotype [*daf-2 (-)*] (Fig. 2D). For the regulation of L2lag, the *daf-18* mutant reduces the duration measured in the wild-type background [*daf-2 (+)*] (wild type vs. *daf-18* p-value=0.0018) and the *daf-2* mutant background [*daf-2 (-)*] (Fig. 2E). In the *daf-2* mutant background, suppression by *daf-18* is stronger than that by *daf-16* (Fig. 2E). Despite this suppression, the *daf-2* mutation increases the duration of L2lag in all backgrounds (Complete statistics in File S1), even in the *daf-16;daf-18* mutant (*t-test* p=0.002) (Fig. 2E), which suggests that DAF-2 also signals to regulate L2lag independently of DAF-18 and DAF-16.

### Mutations in heterochronic genes impact L2lag

L2lag represents a novel feature of the timing of development. To investigate the relationship of L2lag with other developmental events we focused on the heterochronic gene pathway, that controls the timing of cell divisions during postembryonic development. We first addressed the effect of heterochronic genes that impact stage-specific larval events. *lin-14* mutants or RNAi knockdown leads to skipping the L1-specific cell fates. At the level of seam cell divisions, this means that the first intermolt to occur after hatching shows the characteristic symmetric cell divisions of the wild-type L2 stage. *lin-28* mutants or RNAi leads to skipping the L2 cell fates (reviewed in [2] (Fig. 3A). We previously tested the effect of these RNAi treatments in the duration of the larval stages and did not observe major alteration on the duration of the stages in the wild type [7]. When we tested the effect of these mutations in L2lag, we observed that *lin-14* RNAi led to a significant reduction compared to the control RNAi, in both the wild type and *daf-2*. This result could suggest that L2lag is also dependent on the L2 symmetric division. However, *lin-28* RNAi treatment which does not affect L1 events but leads to skipping L2 specific cell fates, did not alter the duration of L2lag (Fig. 3B). These results suggest that L2lag is not related to the L2-specific symmetric seam cell divisions, but it might be a consequence of L1 specific events.

**Figure 3.**
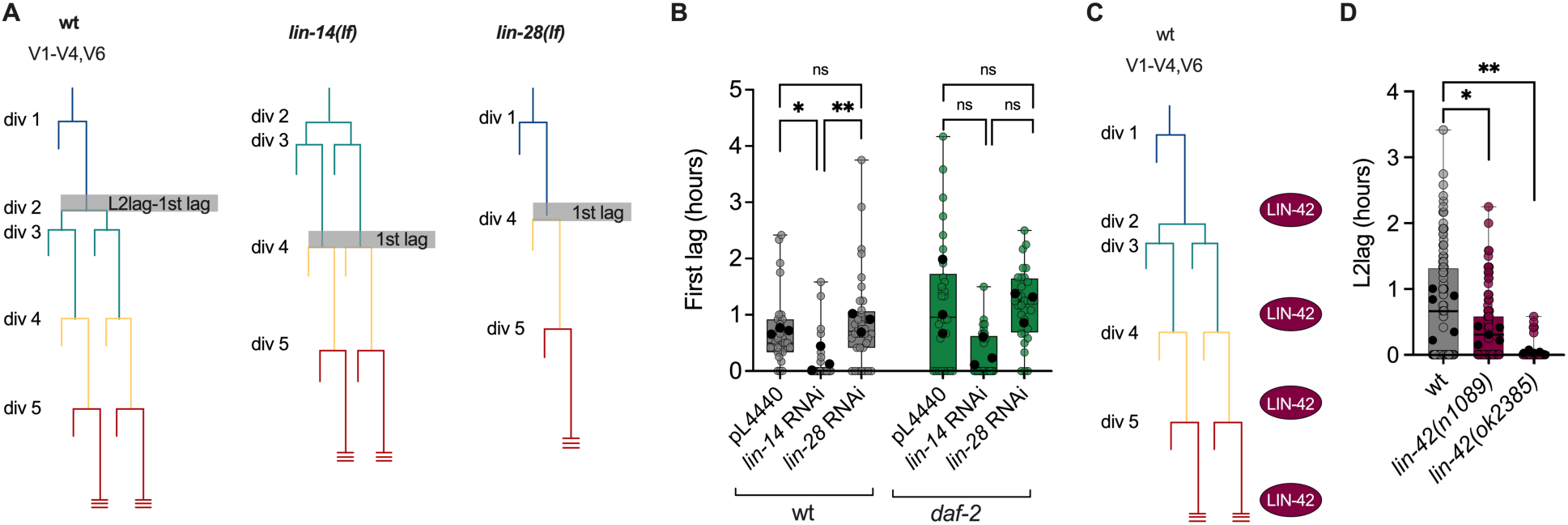
The duration of L2lag is regulated by genes of the heterochronic pathway. **(A)** Diagram of seam cell divisions in the wild - type strain and *lin-14* and *lin-28* lack - of - function mutants. **(B)** Values of the first lag in the wild-type strain and the *daf-2 (e1370)* mutant after treatment with control bacteria (pL4440) or *lin-14* and *lin-28* RNAi. **(C)** Diagram showing the moment of action of LIN-42. **(D)** Duration of L2lag for the wild type and the *lin-42* mutant alleles *n1089* and *ok2385*. For B and D, we performed One-way ANOVA followed by Dunnett’s multiple comparison between the mutants and the wild type, using the experimental averages, * p<0.05, ** p<0.01. Complete report of replicates and statistics results is shown in S2 Table and S1 File. Colored or grey circles show values for individual larvae and black dots show the average of each experiment.

Next, we investigated the impact of another heterochronic gene, *lin-42*, in L2lag. *lin-42* mutants show precocious terminal differentiation [22]. However, unlike *lin-14* and *lin-28*, *lin-42* is expressed periodically during larval molts and its function impacts all larval stages regulating the reiterative process of molting [23–25] (Fig. 3C). Two *lin-42* mutant alleles reduced the duration of the L2lag, but the mutation in *ok2385* had a stronger effect than *n1089* (Fig. 3D). In summary, our results show that the duration of L2lag is impacted by genes of the heterochronic pathways that control the timing of seam cell divisions and molting.

### *daf-2(e1370)* increases desynchrony between ecdysis and seam cell divisions

To investigate relationship between the timing of both molting and cell divisions in the *daf-2* mutants, we focused on seam cells. Seam cells divide asymmetrically at the beginning of each larval stage, and show an additional symmetric division at the L2 stage, prior to the asymmetric one [1]. Specifically, we monitored larvae from hatching to the second larval stage. We performed time-lapse microscopy of single larvae contained in 275×275×30 μm hydrogel microchambers [17]. This setup allows direct observation of hatching and ecdysis as markers of the molting process. Using strains with the reporter *matIs38,* we monitored seam cell divisions of the V lineage. First, we measured the timing of the first two seam cell divisions, hatching and the first ecdysis (Fig. 4A). We analyzed these markers in the wild type, the *daf-2(e1370)* mutant, and the double mutant *daf-2;daf-18*. The first seam cell division (div 1) took place at a similar time relative to hatching for the wild type and *daf-2* but the following event, ecdysis 1, was delayed in the *daf-2* mutant (Fig. 4B). In the wild type, the second seam cell division (div 2) occurred mostly around ecdysis for V1-V4, but V6 usually divides after ecdysis. Div 2 is also delayed in the *daf-2* mutant, but to a larger extent than ecdysis 1. The differential impact of the *daf-2* mutation on the different events creates relative desynchronization between ecdysis 1 and div 2 (Fig. 4D). The *daf-18* mutant showed a level of desynchrony similar to that of the wild type (Fig. S3A). However, in the *daf-2* background, the *daf-18* mutation accelerated divisions so that, in the *daf-2;daf-18* mutant, most divisions occur before ecdysis (Fig. 4B,D). The average timing of the last divisions (V6 lineage) in the wild type occurred at 0.65 hours after ecdysis. For the *daf-2* mutant, this event happened 2.55 hours after ecdysis (*t*-test; p<0.0001) and for the *daf-2;daf-18* mutant at 0.22 hours after ecdysis (*t*-test; p=0.04). These results show that reduced insulin signaling impacts the molting and cell divisions differently, leading to a desynchronization between them. This desynchronization occurs around the transition from L1 to L2, the moment when L2lag takes place. To understand whether seam cell delay (relative to ecdysis) could be related to the L2lag phenotype, we compared their durations across genotypes, observing a statistically significant correlation (Fig. 4E). We also performed the same analysis for the following larval transition, that is, at the end of L2 when ecd 2 and div 4 take place (Fig. 4C). Overall, we observe the same effect of the *daf-2* and *daf-18* mutations, but the desynchrony is less pronounced than at the previous stage (Fig. 4F), consistently with the overall low levels of L3lag (Fig. S3B).

**Figure 4.**
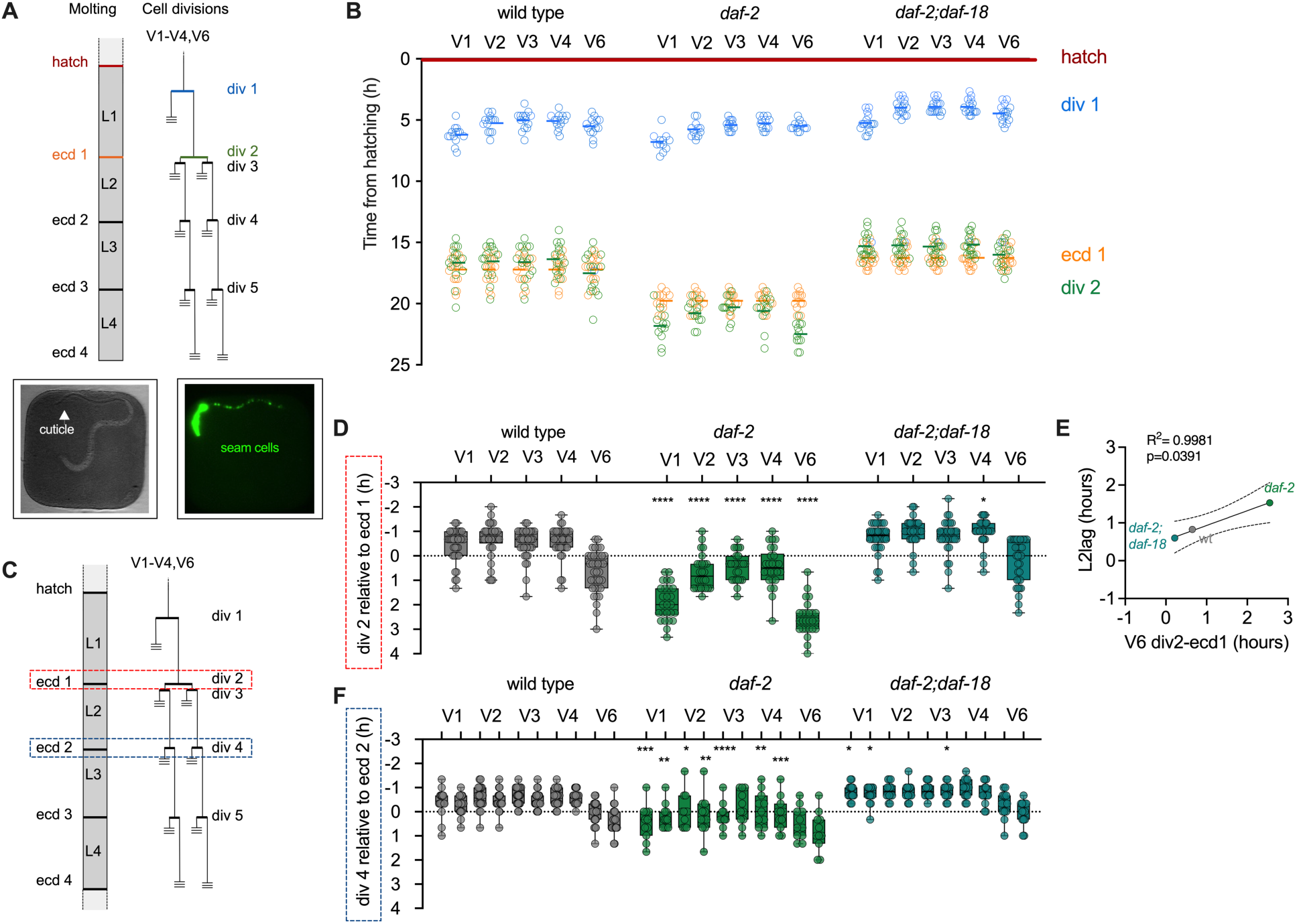
Reduced insulin signaling impacts the two developmental events differently. **(A)** Diagram of the events of the molting process and seam cell divisions larval development, with indication of the events monitored in (B). For seam cell division timing we consider the initiation of divisions. **(B)** Timing of the different developmental events div 1, ecd 1 and div 2 relative to hatching, for each of the seam cells V1-V4, V6. **(C)** Diagram of the events of the molting process and seam cell divisions larval development, with indication of the events monitored in (D) and (F). **(D)** Timing of seam cell division (div 2) relative to ecd 1 for individual larvae of the wild type, the *daf-2(e1370)* mutant and the double mutant *daf-2;daf-18*. **(E)** Correlation between L2lag and V6 seam cell division timing relative to ecdysis (div 2-ecd 1) for the same strains analyzed in B and D. **(F)** Timing of seam cell division (div 4) relative to ecd 2 for individual larvae of the wild type, the *daf-2(e1370)* mutant and the double mutant *daf-2;daf-18*. For D to G, we performed One-way ANOVA followed by Dunnett’s multiple comparison between each of the mutants and the wild type. * p<0.05, ** p<0.01, *** p<0.001, **** p<0.001. Colored or grey circles show values for individual larvae. Complete report of replicates and statistics results is shown in S2 Table and S1 File.

In these experiments, we could observe pumping after ecdysis, independently of the state of seam cell division. Furthermore, we measured the length of the larvae before and after ecdysis, while monitoring seam cell division, and observed that larvae grew at a constant rate after ecdysis, independently of the timing of cell divisions (Fig. S3C). For the same group of larvae, the growth rate was similar between ecdysis and div 2, and after div 2 (Fig. S3D).

Signaling from DAF-2, which is modulated by DAF-18, converges on the transcription factor DAF-16/FOXO, which contributes to cell arrest upon nutrient removal during development [16]. We investigated whether DAF-16 also plays a role in delayed cell division of the *daf-2(e1370)* mutants in food replete conditions. We assessed activation of DAF-16 by analyzing the subcellular localization of DAF-16::GFP in the wild-type and *daf-2(e1370)* mutant backgrounds around the first molt. While the wild-type background showed mainly cytoplasmic localization of DAF-16, the *daf-2(e1370)* mutant showed intermediate/nuclear localization (Fig. S2E). To link DAF-16 activation in *daf-2(e1370)* to delayed seam cell division in fed conditions, we performed time-lapse experiments with the wild type, *daf-2* and *daf-2;daf-16* mutants. The *daf-16* mutation accelerates seam cell divisions of *daf-2(e1370)* similarly to their impact on L2lag (Fig. S2F,G). However, the acceleration from the *daf-16* mutation was not as that from *daf-18*, as the average timing of the last divisions (V6 lineage) of the *daf-2;daf-16* (0.35 hours) was close to that of the wild type (0.12 hours, t-test p=0.3). In summary, *daf-16* and *daf-18* mutations reduce desynchrony between ecdysis and seam cell divisions of the *daf-2(e1370)* similarly to their effect on L2lag and other *daf-2* phenotypes.

We then analyzed another cellular process that takes place around the end of the L1 stage, endomitosis of intestinal cells. During endomitosis, cell enter the M phase of the cell cycle but do not undergo cytokinesis. Endomitosis occurs during L1 in 14 of the 20 intestinal cells, produce binucleated cells [26]. The mutation *daf-2* does not delay the timing of this event relative to ecdysis (Fig. S2J), suggesting that this mutation only impact the timing of some cellular events.

### Desynchrony between developmental events results in L2lag

L2lag and the desynchrony between cell divisions and molting coincide in time, and their magnitude correlates at the population levels when comparing genotypes. Next, we investigated whether the two processes correlate in individual larvae by analyzing the state of seam cells during L2lag. We combined the reporter used for quantification of development by luminometry with a reporter for seam cell division (*matIs38*). We then initiated development of *daf-2* mutants in a luminometer, followed the luminescence profile in real time until the end of the first molt, and collected larvae from three groups. The first group showed a luminometry profile corresponding to the L2lag, with decreasing signal after ecdysis. The second group showed a profile corresponding to a regular intermolt, in this case I2, as determined by a stable, high signal after ecdysis. The third group showed a profile corresponding to a regular intermolt after a period of L2lag (Fig. 5A). For each of the larvae, we then scored the division of seam cells V1-V4 and V6. The L2lag group showed a significantly lower fraction of larvae with divided seam cells than the I2 groups, regardless of whether they had previously passed through L2lag (Fig. 5A). Specifically, only 1 of 9 larvae of the L2lag category had completed all 10 seam cell divisions (V1-V4, V6; left and right). This result suggests that L2lag could indeed be a consequence of the desynchrony between the molting and cell division processes. Additionally, we measured luminescence over the L1 to L2 transition in a microscope, coupled to imaging of both ecdysis and seam cell divisions. We confirmed that ecdysis always occurs linked to the increase of signal after approximately two hours of reduced luminescence signal. Then, larvae that reached ecdysis before completing seam cell division showed in general lower levels of luminescence (Fig. S4). When the different categories of larvae were averaged, there was a clear difference in bioluminescence between animals that had already finished seam cell divisions and those that did not. In particular, the larvae that had not completed divisions before ecdysis showed a reduction in the signal that, after approximately 2.5 hours reached again the levels of luminescence measured after ecdysis (Fig. 5B). This result suggests that delayed developmental events, evidenced by seam cell divisions, lead to entry into L2lag.

**Figure 5.**
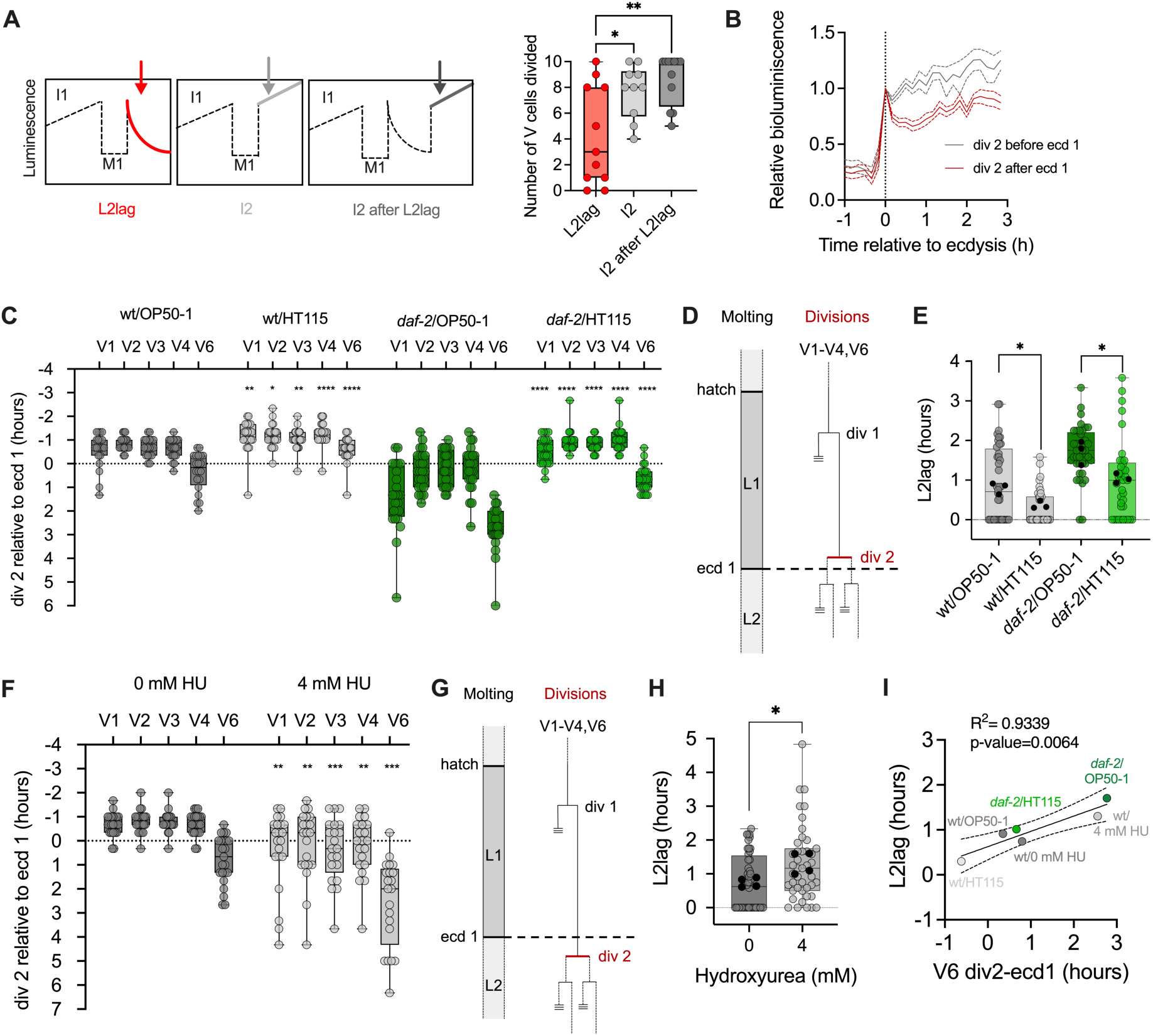
Modification of seam cell division timing relative to ecdysis alters the duration of L2lag. **(A)** Diagram showing the moment of collection (arrows) of the three categories of larva according to their luminometry profile (left) and fraction of animal showing divided seam cells (V1-V4, V6) for each of the three categories (right). The fraction of animals was calculated from 9 larvae in L2lag, 10 larvae in I2, and 9 larvae in I2 after L2lag. We performed two-way ANOVA followed by Tukeýs multiple comparison to assess the effect of the group across all cells analyzed. **(B)** Average bioluminescence relative to ecdysis (time=0), for larva that show divisions before (14 wt larvae) or after (18 *daf-2* and 2 wt larvae) ecdysis 1. The signal from each larva is normalized to the bioluminescence level at ecdysis (bioluminescence=1 at t=0). Continuous lines represent the average bioluminescence value. Dashed lines represent s.e.m. **(C)** Analysis of div 2 relative to ecd 1 in the wild type and the *daf-2* mutant fed with OP50-1 and HT115. **(D)** Diagram showing the situation of div 2 relative to ecdysis favored by feeding with HT115. **(E)** Duration of L2lag in the wild type and *daf-2(e1370)* fed with OP50-1 and HT115. **(F)** Analysis of div 2 relative to ecd 1 in the wild type treated with 4 mM Hydroxyurea, and without treatment. Statistics analysis reflect results from unpaired t-test comparing the timing of each division between the two conditions. **(G)** Diagram showing the situation of div 2 relative to ecdysis favored by the treatment with Hydroxyurea. **(H)** Duration of L2lag for larvae treated with 4 mM Hydroxyurea, and without treatment, in four independent experiments. We performed One-way ANOVA followed by Dunnett’s multiple comparisons to test differences between each of the treatments and the untreated larvae. Colored and grey circles show values for individual larvae and black dots show the average of each experiment. For the data on panels A and I, we performed One-way ANOVA followed by Tukeýs multiple comparisons to test differences between all strains. For the data on panels C, E, F and H, we performed unpaired t-test to compare the effect of the diet within each strain or that of HU. Complete report of replicates and statistics results is shown in S2 Table and S1 File. Statistically significant differences are marked as * p<0.05, ** p<0.01, ***p<0.001, **** p<0.0001.

To test this hypothesis, we experimentally tested conditions that modified the timing of seam cell divisions relative to ecdysis. We expected that advancing seam cells divisions would reduce L2lag and delaying seam cell divisions would increase it. We first advanced seam cell divisions relative to ecdysis by growing the larvae on *E. coli* HT115, typically used for RNAi. HT115 accelerate the timing of postembryonic development [27], but its role on seam cell division has not been assessed. While feeding with HT115 might impact additional processes, we could establish that HT115 significantly advanced seam cells divisions compared to OP50-1, in both the wild type and *daf-2(e1370)* (Fig. 5C), without impacting the timing of ecdysis (Fig. S5A). This led to a situation where most divisions occur before ecdysis (Fig. 5D). Indeed, when we measure L2lag, we found a significant reduction in both the wild type and *daf-2(e1370)* (Fig. 5E), again, without altering the duration of L1 measured by luminometry (Fig. S5B).

We also tried to delay the cell divisions by treating developing larvae with Hydroxyurea (HU), which prevents DNA synthesis by replication fork arrest. Since the delay in divisions could also feedback to the molting process, we aimed for concentrations of Hydroxyurea that allowed complete development and did not increase the duration of the first larval stage. Hydroxyurea at 4 and 8 mM allowed progression through postembryonic development (Fig. S5C), and only 4 mM permitted developmental progression without significant delay of the first larval stage (Fig. S5D). As before, treatment with HU might impact other processes. To confirm the impact of HU on the timing of seam cell divisions relative to ecdysis, we followed these two features on the larvae treated with 4 mM HU. The treatment with HU significantly delays divisions relative to 0 mM HU (Fig. 5F), without impacting the timing of ecdysis (Fig. S5E). This provokes that many cell divisions occur after ecdysis (Fig. 5F-G). Indeed, at 4 mM Hydroxyurea the L2lag was significantly prolonged compared to the control condition (Fig. 5H). For both HT115 feeding and treatment with HU, we plotted the L2lag duration against the timing of seam cell division, finding a significant positive correlation (Fig. 5I, Fig. S5F). These results show that varying the relative desynchrony between ecdysis and cell divisions modulates the duration of L2lag.

Interestingly, this also explains our findings on how *lin-42(ok2385)* mutation impacts L2lag. Previous research showed that the *lin-42(ok2385)* increased the duration of the L1 stage but showed a timing of seam cell divisions similar to the wild-type, resulting in advanced cell divisions relative to ecdysis [23,24], which is consistent with the reduced L2lag we find in this mutant (Fig. 3D). This suggests that the effect of LIN-42 on L2lag is likely related to its molting clock function [25].

Finally, we asked whether increased L2lag is a general principle for other interventions that affect the duration of development. *C. elegans* development shows a strong temperature dependence [28]. We focused on this environmental cue as, despite the slowed developmental rate at lower temperature, the events of cell divisions and ecdysis occurred at the same time relative to the duration of the total development [6]. This means that reduction of speed due to temperature does not involve increased desynchrony between ecdysis and cell divisions. Since we previously characterized the regulation by temperature of postembryonic development for temperature between 10 and 27 °C [7], we could extract information about L2lag from that dataset. Despite the change in developmental rate, L2lag showed similar values at all temperatures tested (Fig. 6A). When analyzed in time-lapse experiment, the L1 stage (hatch to ecd1) was about 3 hours faster at 22.5 °C than at 20 °C (Fig. 6B) but the timing of seam cell division relative to ecd 1 was similar between conditions (Fig. 6C), consistent with the lack of differences in L2lag. Likewise, L2lag is similar between 12 and 22 °C for the *daf-2* mutants (Fig. 6D), as despite of acceleration of development (Fig. 6E), temperature did not impact overall seam division timing (Fig. 6F). This means that seam cell divisions are not *per se* more susceptible to environmental interventions that reduce developmental rate. Additionally, culture of *daf-2(e1370)* mutants at 22 °C leads to dauer formation, an alternative state to L3 stage. The similar levels of L2lag in the *daf-2* mutant at 20 °C and 22 °C, as well as the similar levels of seam cell desynchrony, confirm that L2lag does not correlate with dauer formation. Consequently, increased L2lag is neither a trivial nor a direct consequence of slow development.

**Figure 6.**
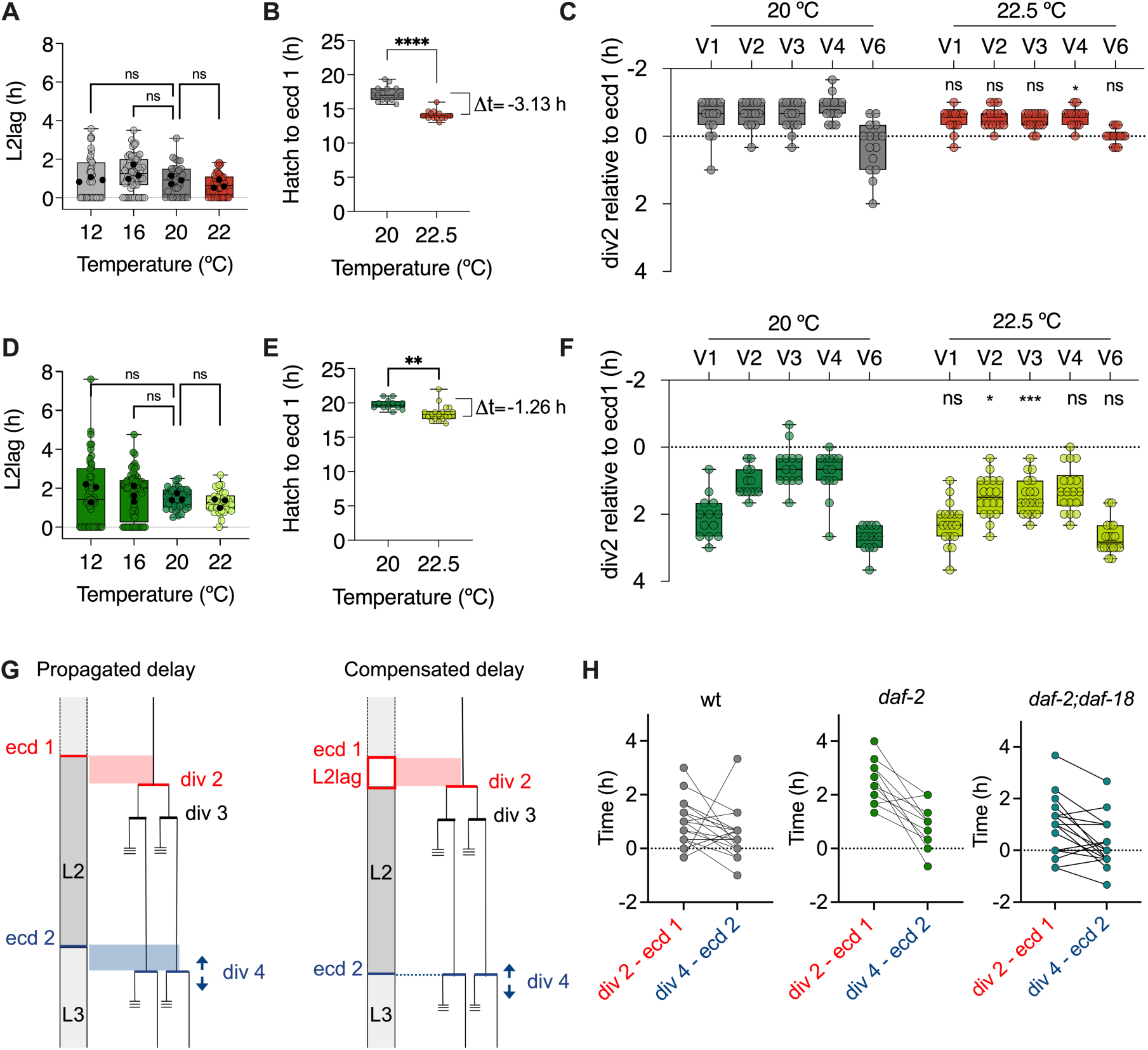
Temperature dependent acceleration of development does not impact the duration of L2lag. **(A)** Duration of the L2lag stage for the wild-type strain at 12, 16, 20 and 22 °C in three independent experiments. **(B)** Duration of the L1 (from hatching to ecdysis 1) recorded in time-lapse imaging experiment for the wild type at 20 °C and 22.5 °C. **(C)** Timing of seam cell division (div 2) relative to ecd 1 for individual larvae of the wild type at 20 °C and 22.5 °C. **(D)** Duration of the L2lag stage for the *daf-2(e1370)* mutant at 12, 16, 20 and 22 °C in three independent experiments. **(E)** Duration of the L1 (from hatching to ecdysis 1) recorded in time-lapse imaging experiment for the *daf-2(e1370)* mutant at 20 °C and 22.5 °C. **(F)** Timing of seam cell division (div 2) relative to ecd 1 for individual larvae of the *daf-2* mutant at 20 °C and 22.5 °C. For A and D, we performed One-way ANOVA followed by Dunnett’s multiple comparison between each of the temperatures and the standard (20 °C). For B, C, E and F, we performed unpaired *t*-test. Complete report of replicates and statistics results is shown in S2 Table and S1 File. Colored or grey circles show values for individual larvae and black dots show the average of each experiment. **(G)** Diagrams showing the effect of propagation or compensation of the delay of div 2 (relative to ecd 1) on the duration of the delay of div 4 (relative to ecd 2). **(H)** Plots showing the measurements of div 2 relative to ecd 1 and div 4 relative to ecd 2 for individual larvae of the wild type, *daf-2* and *daf-2;daf-18* mutants.

Taken together, our results support the idea that the initiation of larval stages can be delayed when events of the previous stage did not complete. As we have observed, the alteration in the order of developmental events is tolerated by the system. However, such a mechanism would prevent that desynchrony provoked by a delayed events amplifies over the course of development.

## DISCUSSION

Using a method for continuous monitoring of *C. elegans* development, we identified a previously unrecognized lag phase between ecdysis and the onset of the following intermolt under fed conditions (Fig. 1). Such transient states are unlikely to be detected by morphological observations alone, underscoring the importance of high-throughput, single-animal analyses to uncover deviations from average developmental timing. While our current assays cannot resolve the mechanisms underlaying reduced luminescence signal during the lags, our results show that these lags arise when stage-specific cellular events did not complete within the boundaries defined by molting. This indicates, even in the wild-type background and in replete conditions, developmental progression can pause at the beginning of intermolts, We propose that this pause could represent a mechanism to accommodate resynchronization between developmental events.

L2lag is more frequent and lasts longer in *daf-2(e1370)* mutants, but this phenotype is independent of other developmental phenotypes of the mutant. First, the duration L2lag does not correlate with that of other stages (Fig. S1). Second, the variability of L2lag is approximately one order of magnitude higher than that of other stages (Fig. 1H). Third, the *daf-16* mutation completely suppresses the extended duration of intermolts but not the increased duration of L2lag (Fig. 2B,C).

Robust development requires coordination between cell divisions and their differentiation, and organismal growth (reviewed in [29]). In *C. elegans*, the timing of molting and cell divisions can be uncoupled, and environmental cues may affect these processes either uniformly or differentially. Because reduced insulin/IGF-1 signaling (IIS) could impact the two processes differently, we investigated cell divisions dynamics. We observed that seam cell divisions are more sensitive than the molting program to reduced IIS (Fig. S2A-C). Our results suggest that IIS independently impacts the two timers acting on *C. elegans* development, those controlling the timing of molting and cell divisions. Although reduced IIS slows both processes, it does so to different degrees, leading to a delay on seam cell division relative to ecdysis (Fig. 4D), that is, a desynchronization between developmental events. This desynchronization is linked the L2lag: *daf-2(e1370)* in L2lag are more likely to have undivided seam cells than larvae at the onset of the second intermolt (Fig. 5A). Thus, the lag phase we observe before the L2 intermolt likely serves as a buffering mechanism that allows delayed seam cell divisions to catch up. Prior work shows that interfering with L2-stage cell divisions causes exposure of a defective L3 cuticle and larval death [5], indicating that excessive delays in cell division relative to molting can be lethal and that mechanisms ensuring resynchronization are essential. Similar checkpoint-based coordination is well described in *Drosophila*, where damaged or slow-growing imaginal discs secrete DILP8, which suppresses ecdysone production and delays metamorphosis until growth is synchronized [30,31]. Similarly, the lag phase of the molting process after the first molt observed in our work delays the initiation of the L2 stage, providing time for slower developmental events to realign with the molting cycle.

Not all interventions that delay development increase L2lag. The temperature dependence of development in *C. elegans* follows the Arrhenius equation, which describes the relationship between temperature and the rate of first-order chemical reactions [7]. At temperatures between 15 and 23 °C, cell divisions and ecdysis occur at the same times relative to the duration of development [6]. With this previous information, we predicted that temperature should not impact L2lag. Indeed, L2lag is similar at temperatures between 12 and 22 °C.

Our investigation offers other insights about *C. elegans* developmental timing. The lag phases occur at the beginning of larval stages, the same moment when a nutritional checkpoint have been described. Starvation causes arrest at different stages: newly hatched larvae arrest in L1 when food is absent [32,33], and starvation during L2 and L3 leads to arrest at the beginning of L3 and L4 respectively [16]. When food is removed mid-stage, larvae progress only to the onset of the next stage, consistent with checkpoint control. L2lag, L3lag and L4 lags may therefore represent delays to pass these nutritional checkpoints. Supporting this idea, fed *daf-2(e1370)* mutants grown to mid-L2 at 15 °C and then shifted to 25 °C paused preferentially early in the L3 and L4 stages [16,34]. If L2lag reflects a delay at this checkpoint, our data suggest that completion of cell divisions may constitute the requirement for checkpoint passage. Notably, another process occurring at stage boundaries is the arrest and re-initiation of molting-coupled gene expression oscillations [4], which begin ∼5 hours after release from L1 arrest—close to the timing of the first seam cell division [1].

Because molting and seam cell divisions are not intrinsically coupled, a delay in div 2 relative to ecd 1 could, in principle, be propagated to the next transition (div 4 relative to ecd 2). Then this delay could be increased or decreased due to the intrinsic variability in speeds within the L2 stage (Fig. 6G). Alternatively, if each larval stage functions as a modular unit gated by a checkpoint, delays in earlier events could be compensated by postponing initiation of the next intermolt (Fig. 6G). As before, the intrinsic variability of the L2 would also increase or decrease the delay of div 4 relative to ecd 2. Our time-lapse data support the second model: across genotypes, larvae showing a delay of div 2 relative to ecd 1 consistently exhibited reduced desynchrony in the next stage (div 4 relative to ecd 2) (Fig. 6H). This indicates that delays do not propagate across stages. Such compensation confines perturbations to a single stage, resembling developmental modularity, which enhances evolvability by allowing independent optimization of stage-specific processes (reviewed[35]). Consistent with this, we previously showed that *C. elegans* larval stages have distinct timings and respond differently to rate-limiting factors [7], further supporting the idea that each stage’s duration reflects the developmental events specific to that stage.

## MATERIALS AND METHODS

### Caenorhabditis elegans

*C. elegans* strains were cultured following standard methods [36]. Animals were grown and maintained on nematode growth media (NGM) with a lawn of *Escherichia coli* OP50-1 at 20 °C. Experimental temperature was 20 °C, except when stated otherwise. All strains used are listed in S1 Table.

### Bacterial strains

For maintenance plates and experiments the food source was *E. coli* OP50-1. The only exceptions are Fig. 3B, and 5C,E, and S5A,B, where we used *E. coli* HT115.

### Luminometry of single worms

To analyze the developmental timing of single worms we used the bioluminescence-based method previously described [11]. To obtain age-synchronized eggs, 10-15 hermaphrodites were transferred to a fresh NGM plate, allowing them to lay eggs for one hour. For experiments with six conditions or less, synchronized-eggs were individually placed in wells of a 96 well-plate containing 100 μl of S-basal (5 μg/ml cholesterol) with 200 μM Luciferin. After placing all embryos, 100 μl of S-basal with 20 g/l *E. coli* OP50-1 or HT115 was added to each well. For experiments with more than 6 conditions, the eggs were placed individually in wells of a 384-well plate containing 25 μl of S-basal (5 μg/ml cholesterol) with 400 μM Luciferin, adding 25 of S-basal with 20 g/l *E. coli* OP50-1 (or HT115 for Fig. 5E and Fig. S5B) per well after all eggs were transferred. In both cases, samples from different strains or conditions were alternated to avoid local effects. Plates were sealed with a gas-permeable membrane and placed in a luminometry reader (Berthold Centro XS3) localized inside a cooled incubator (Panasonic MIR-254). Luminescence was recorded for 1 s (for 96-well plate) or 0.7 s (for 384-well plate) at 5-min intervals until reaching adulthood.

### Temperature regulation

We measured the temperature in the plate using a data logger Thermochron iButton DS1921G (Maxim Integrated). The temperature in the plate is ∼3.5 °C higher than that set at the incubator, due to the production of heat from the luminometer. All temperatures shown in the data correspond to the temperature experienced by the larvae.

### Live time-lapse imaging of single worms

To study gene expression, progression of molting and seam cells divisions of single worms we adapted a system for time-lapse microscopy in [17]. To prepare the acrylamide-gel chambers, we used micropatterned silicon wafer with microchambers dimensions of 275×275×30 μm. The microchambers were prepared by first mixing a polyacrylamide solution containing 10 % acrylamide/bis-acrylamide (29:1), 1 % ammonium persulfate and 0.1 % TEMED. The liquid solution was pipetted into a compartment composed by the master wafer glued to a methacrylate frame. After filling all the compartment, a glass microscope slide was placed on top to seal the compartment and let it to solidify for 0.5-2 hours. The resulting polyacrylamide gel is washed with ddH_2_O at least 4 times and stored in ddH_2_O.

To perform the experiments, the polyacrylamide microchamber used is previously stabilized in M9 during 1 hour prior to the experiment. To prepare the long-term imaging, a methacrylate frame is stuck to a microscope slide. The M9-stabilized microchamber is then placed on top of the slide, within the frame. To place embryos in the polyacrylamide chambers we proceeded as previously described [37]. We obtained age-synchronized embryos by transferring 10-15 gravid hermaphrodites to fresh NGM plates and letting them to lay eggs for one hour. Using an eyelash, eggs were individually transferred to the wells of the microchamber (1 egg per well). After eggs were transferred, we collected some OP50-1 from NGM plates using the eyelash and added it to wells as food source. The only exception is for Fig. 5C and S5A, that we used OP50-1 or HT115. The compartment was filled with M9 and sealed with cover slides.

We performed the time-lapse imaging in a Leica scope M205 FCA with a motorized stage, using GFP excitation/emission filters. Time-lapse was performed by taking pictures every 20 minutes for each worm in a Z-stack of 40 μm divided in 11 steps. Worms were photographed in two different channels (white light channel and GFP channel) with exposure times of 50 ms for both cases. The developing larvae worms were recorded for approximately 40 hours. For figure S3F, we used strains that contains the reporter *sevIs1* in addition to *matIs38.* Since *sevIs1* expresses GFP ubiquitously, we visualized seam cells using the RFP signal from *matIs38.* For this experiment we used the Leica M205 FCA scope and a Nikon Eclipse Ti2 microscope.

### Analysis of time-lapse microscopy

For the analysis of expression of *dpy-9*p::GFP expression we selected the best-focused Z step for every timepoint (GFP channel). Using ImageJ software, we calculated the expression by obtaining the average GFP intensity of 3 middle-body cells after subtracting the background. We normalized the GFP intensity levels to the average values of first three hours before ecdysis. For Fig. 1J, we smoothed the signal by applying a centered moving average over 1 hour (5 time points). For the analysis of molting progression, we reported hatching when found any part of the larvae outside of the eggshell. Likewise, ecdysis is marked when the larvae partially or completely exited the old cuticle.

For the analysis of seam cell divisions, we reported each specific seam cell (V1-V4, V6) as divided when at least one of the sides presented evidence of cytokinesis. For the analysis of intestinal cell endomitosis, we reported the timing of initiation of nuclear divisions. Unlike seam cell division, intestinal cell endomitosis occurs with high synchrony throughout the larva.

### Seam cell division analysis at specific larval stages

We analyzed the state of seam cell division specific developmental stages. To do so, we perform the luminometry assay as described above (see “*Luminometry of single worms*”) using worms combining the bioluminescence (*sevIs1*) and the seam cell fluorescence-tagged reporter (*matIs38*). After approximately 20 hours of initiating, we monitored the larval developmental progress continuously. At a single timepoint, we collected larvae that were in the different categories to be analyzed. During each replicate, we extracted only 4-5 larvae from the complete 96-well plate to maintain all the processing time of the samples lower than 30 min. We collected animals that, at the time of collection, were within 0.5-1 hour from the previous transition, that is, from the end of M1 for animals in the categories “L2lag” and “I2”, and from the end of L2lag for the larvae in the category “I2 after L2lag”. Individual worms were isolated and washed from food M9. Finally, worms were immobilized using 10 mM Levamisol diluted in M9 and placed in agarose pads. Larvae with seam cell divisions were identified using a Leica scope M205 FCA and RFP excitation/emission filters.

### Quantification of GFP and movement at specific larval stages

To collect larvae in the different stages, we proceeded as explained above. Then, for quantification of GFP, we place individual larvae in a drop of approximately 10 μl of S-basal on a microcopy slide and added 1 μl of 10 mM Levamisol. We acquired pictures in using a Leica scope M205 FCA and GFP excitation/emission filters. We manually segmented the complete larvae and measure average pixel intensity with Image J. For the quantification of movement, we placed the larvae on NGM plates, measured the tracks left by the movement of the larvae after approximately 30 min, and then normalized to the exact time for each larva.

### Luminescence recording by time lapse imaging

For luminescence measurement in microchambers, we prepared the acrylamide chambers as previously reported but the gels are stabilized in M9 containing 2.5 mM Luciferin. After placing the eggs, the compartment is filed with the same solution. The system is transferred to a microscope Nikon Ti2 Eclipse equipped with a camera for transmitted light and RFP (Nikon DS-Qi2) and an additional camera with high-sensitivity CCD sensors (iKon-M 934 CCD; Andor). For bioluminescence, we integrated the signal form 1-min exposure. To obtain the average bioluminescence signal we use the ImageJ software. As larvae move over the 1-min exposure, we draw a region of interest (ROI) that frames the entire well to integrate the signal from the complete acquisition time, then obtaining the average bioluminescence value per pixel. After this, we subtracted the background calculated by drawing a ROI where the worm is absent, then obtaining the average background value. In this experiment (Fig. 5B), we used strains that contains the reporter *sevIs1* for luminescence and *matIs38* to track seam cell divisions. Since *sevIs1* expresses GFP ubiquitously, we visualized seam cells using the RFP signal from *matIs38.* For white-light and RFP we acquired in a Z-stack of 44 μm divided in 11 steps. For white-light, exposure time was 70 ms and for RFP, exposure time was 200 ms.

### Hydroxyurea treatment

To test the effect of cell division blocking chemicals on the development, we treated worms with Hydroxyurea (HU) specifically during the postembryonic development. To do so, we performed the luminometry assay as described above (see “*Luminometry of single worms*”). The difference is that eggs were placed in wells containing 90 μl of S-basal (5 μg/ml cholesterol) with 200 μM Luciferin and after placing them, we added 90 μl of S-basal with 20 g/l *E. coli* OP50-1 to each well, so the final volume is 180 μl per well. 12 hours after the beginning of the luminometry recording, the plate was unloaded from the luminometer. For non-treated, we added 20 μl ddH_2_O per well. For the 4 mM HU treatment, 40 μl of the 1 M HU stock was diluted in 960 μl ddH_2_O and 20 μl of that mix was added to each well. For the 8 mM HU treatment, 80 μl of the 1 M HU stock was diluted in 920 μl ddH_2_O and 20 μl of that mix was added to each well. After adding the HU, the plate was sealed with a new gas-permeable membrane and placed again in a luminometry reader.

For HU treatment in microchambers, we proceeded as previously described for time-lapse imaging, but non-treated worms and 4 mM HU treated worms were placed in separated. The two gels were placed on the same slide and imaged together. For the 4 mM treatment, the gel was stabilized in M9 with 4 mM HU, and after placing the eggs, the compartment was filled up with the same solution.

### Statistical analysis

For statistics we have used the averages of independent biological replicates to avoid the inflated N value from using individual animals in luminometry experiments [38]. Before calculating the averages, we applied a filter for outliers using ROUT (Q=0.2%) on Graphpad Prism 10. For Fig. 3D, we used iterative Grubbś, as ROUT removed all the values different from The number of larvae and number of independent replicates performed are shown in Table S2. For the data on Fig. 1G we used two-way ANOVA to test both the effects of genotype and larval stage, followed by Fisheŕs LSD to reveal significant differences between genotypes at each stage. We used unpaired two-tailed t-test to compare the means of two groups in Fig. 5C, E, F, H, Fig. S5A, B, E and Fig. 6B, C, E and F. For the rest of the experiments, we used one-way ANOVA to compare more than two groups. After one-way ANOVA, we either perform Dunnett’s multiple comparison to compare the average of each genotype/condition to the average of the wild type, or Tukeýs multiple comparison to compare the average of each genotype to all the other genotypes. The type of analysis that applied to each panel is detailed in the figure legends. For clarity, some plots show only relevant comparisons, but all the results from statistical testing are shown in S1 File. In all cases, * means p>0.05, ** p>0.01, *** p>0.001, and **** p>0.0001. Graphs and statistics were performed on Graphpad Prism 10.

## ACKNOWLEDGEMENTS

Some strains were provided by the *Caenorhabditis* Genetics Center (CGC), which is funded by NIH Office of Research Infrastructure Programs (P40 OD010440). Fig. 2A was created with <u>BioRender.com</u>. Work in the Olmedo lab is supported by the grants PID2022-139009OB-I00 funded by MICIU/AEI/ 10.13039/501100011033 and by “ERDF A way of making Europe”, and P20-RT-01248 (PAIDI2020, Proyecto cofinanciado por fondos del Programa Operativo FEDER Andalucía 2014-2020 y por la Consejería de Transformación Económica, Industria, Conocimiento y Universidades de la Junta de Andalucía). Initial observations of *daf-2* delayed divisions were performed by M.O. at the lab of J.v.Z during a COST GENiE Short Term Scientific Mission granted to M.O. (COST-STSM-BM1408-35915). FJRE was supported by a contract of the VPPI-University of Sevilla (Contratos de Personal Investigador en Formación), and a FEBS short-term fellowship. MMB was supported by a postdoctoral contract from the “Programa de Ayudas a la I+D+i, en Régimen de Concurrencia Competitiva en el Ámbito del Plan Andaluz de Investigación, Desarrollo e Innovación (PAIDI 2020)”. AMC was supported by a contract of the VII PPIT- University of Sevilla (Contratos de Acceso al Sistema Español de Ciencia, Tecnología e Innovación). The Nikon Eclipse Ti2 microscope system was funded by DFG (INST 86/1851-1 FUGG). The funders had no role in study design, data collection and analysis, decision to publish, or preparation of the manuscript.

## AUTHOR CONTRIBUTIONS

Conceptualization: FJRE, AMC, MO

Formal analysis: FJRE, AMC, MO

Investigation: FJRE, AMR, MMB

Methodology: FJRE, FS, NG, MM, JvZ, MO

Visualization: FJRE, MO

Supervision: AMC, MO

Writing—original draft: FJRE, AMC, MO

Writing—review & editing: FJRE, AMR, MMB, FS, NG, MM, JvZ, AMC, MO

## SUPPLEMENTARY FIGURES

**Figure S1. (Relative to figure 1).**
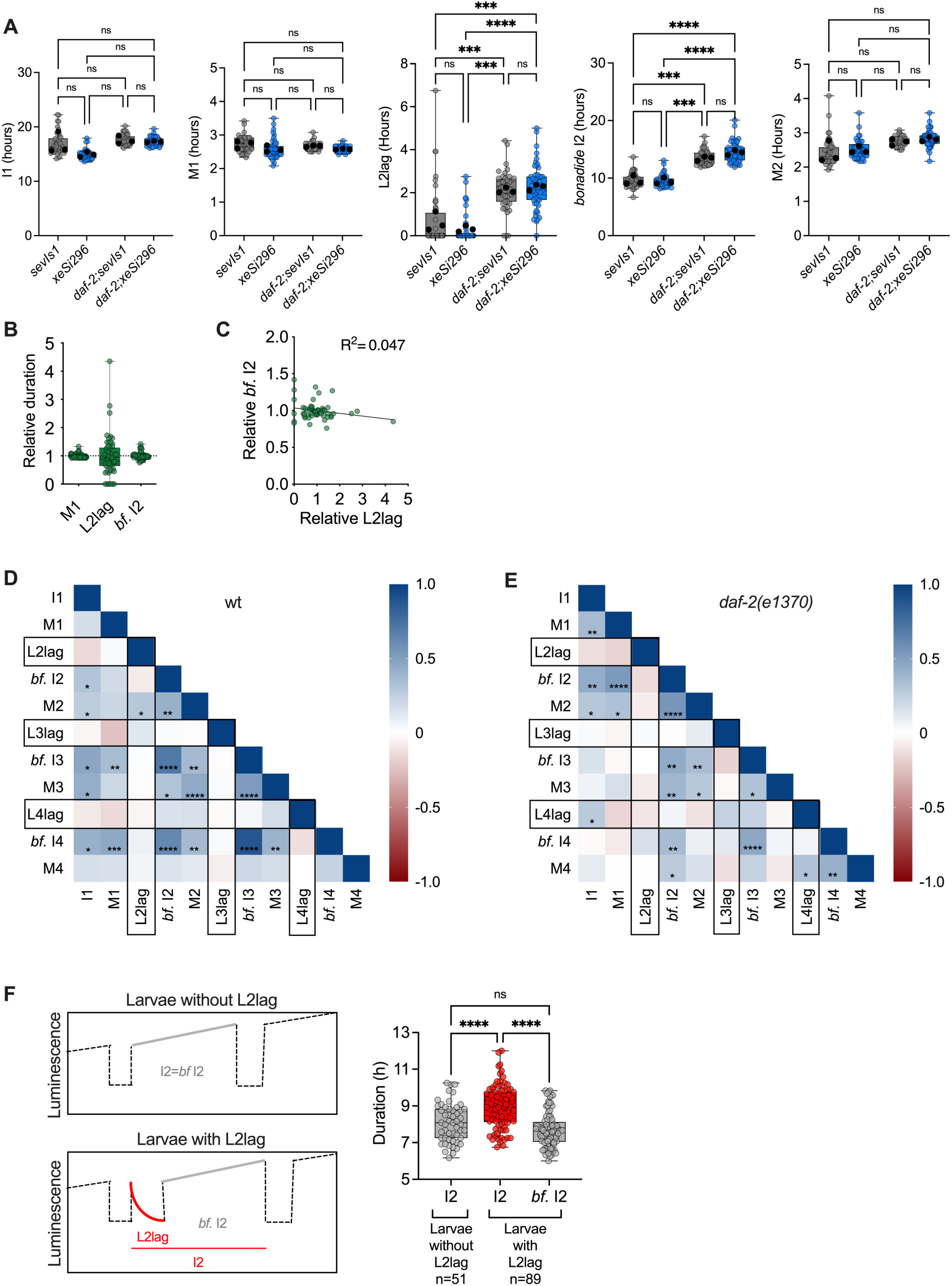
**(A)** Duration of I1, M1, L2Llag, bonafide I2 and M2 for wild type and *daf-2* mutant with the luminescence reporters *sevIs1* and *xeSi296*. **(B)** Duration of M1, L2lag and *bonafide* I2 relative to the average duration of each stage for the same larvae shown in F. **(C)** Relative duration of L2lag and bf. I2 for each of the larvae in F and H.**(D-E)** Matrix heatmap of the correlation between the duration of all larval stages in 58 individual larvae of wildtype (D) and 57 individual *daf-2(e1370)* mutant larvae (E). **(F)** Diagrams showing the luminescence profile and durations analyzed, for wild-type larvae without and with L2lag (left), and duration of the corresponding periods (right).

**Figure S2 (Relative to figure 2).**
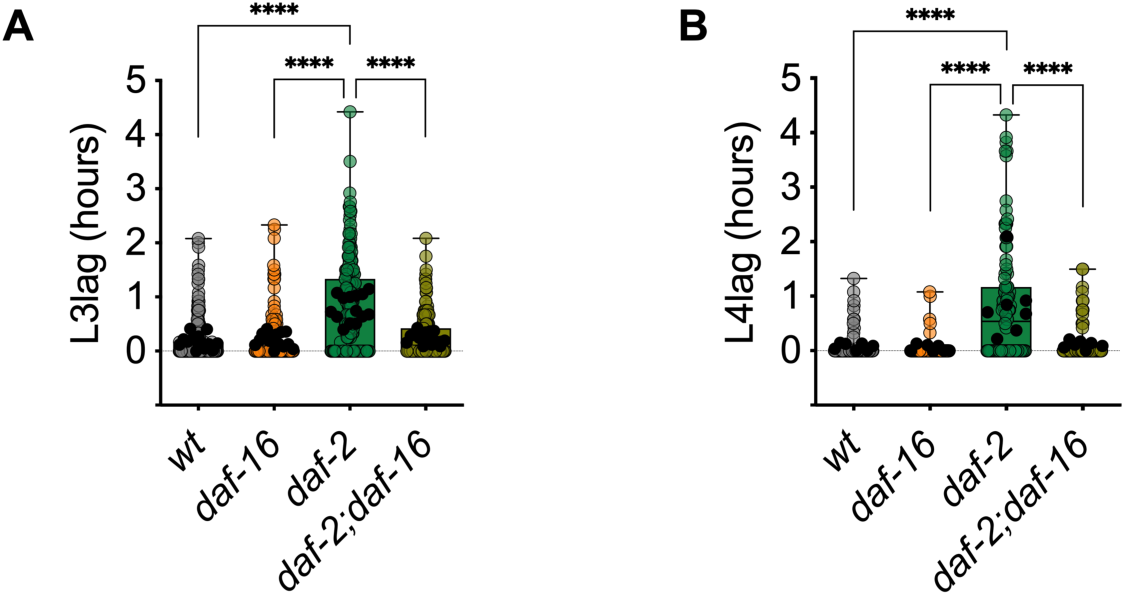
Duration of L3lag **(A)** and L4lag **(B)** for the wild type, *daf-2*, *daf-16,* and *daf-2;daf-16*. Colored circles show values for individual larvae and black dots show the average for each experiment. For both panels, we performed One-way ANOVA followed by Tukeýs multiple comparisons to test differences between all strains. Only significant differences are shown, **** p<0.0001.

**Figure S3 (Relative to figure 4).**
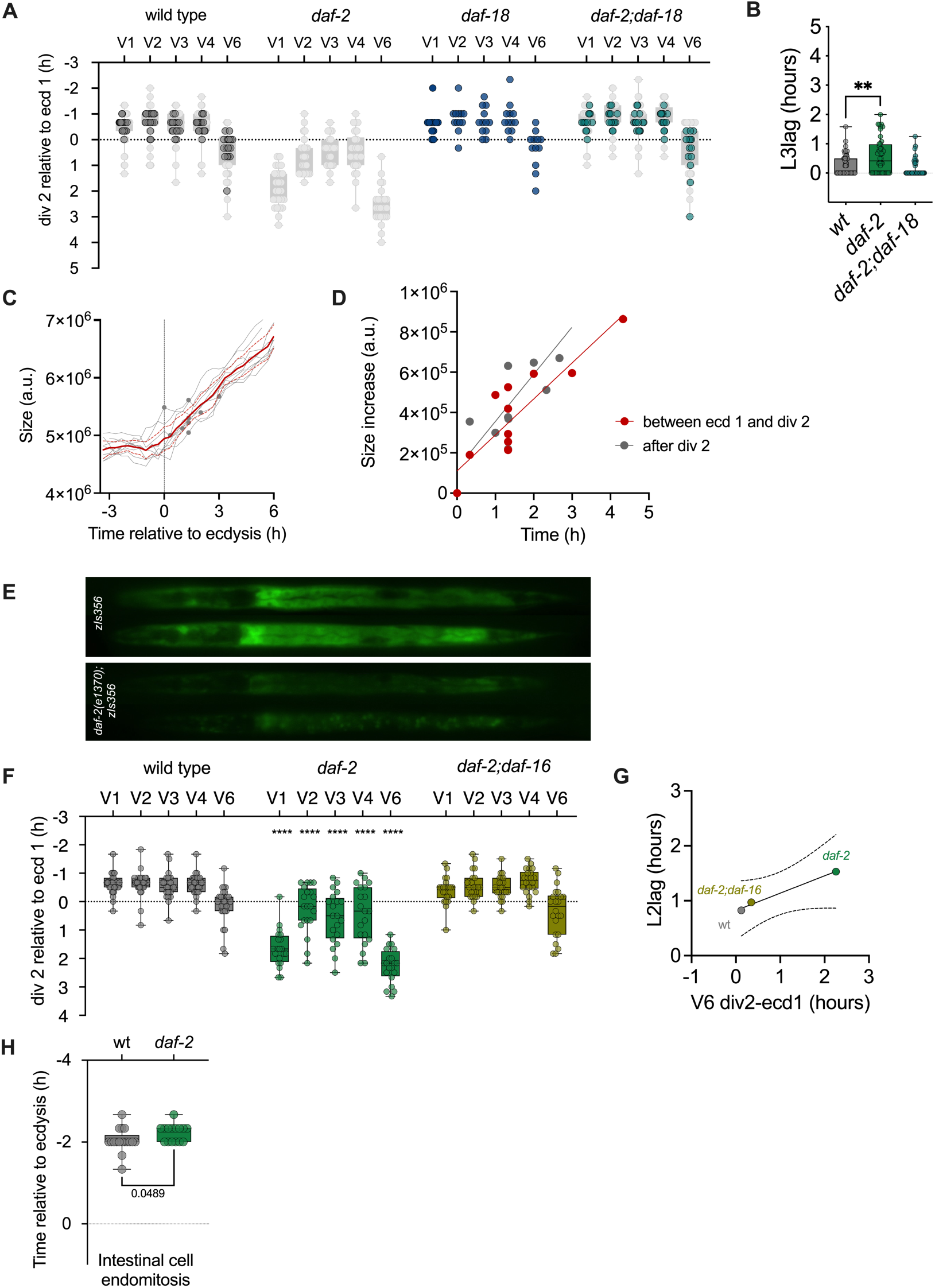
**(A)** Timing of seam cell division (div 2) relative to ecdysis (ecd 1) for the wild type, *daf-18* and *daf-2;daf-18*. This data set is shown overlaid to the data presented in Fig. 4D. **(B)** Duration of L3lag for the wild type, *daf-2* and *daf-2;daf-18*. **(C)** Quantification of size relative to the timing of ecdysis. Gray lines show the values of individual larvae and red line shows the average (and 95% CI). Gray dots show the timing of seam cell division for each animal. **(D)** Size increase per time interval between ecd 1 and div 2 (red) and in a similar period of time after div 2. **(E)** Representative images of DAF-16::GFP on the wild type and *daf-2(e1370)* backgrounds. **(F)** Timing of seam cell division (div 2) relative to ecd 1 for the wild type, the *daf-2* mutant, and the double mutant *daf-2;daf-16*. Colored circles show values for individual larvae. **(G)** Correlation between L2lag and the timing of V6 seam cell division relative to ecdysis (div 2-ecd 1). **(H)** Timing of intestinal cell endoreplication relative to ecdysis, for the wild type and *daf-2*.

**Figure S4. (Relative to figure 5).**
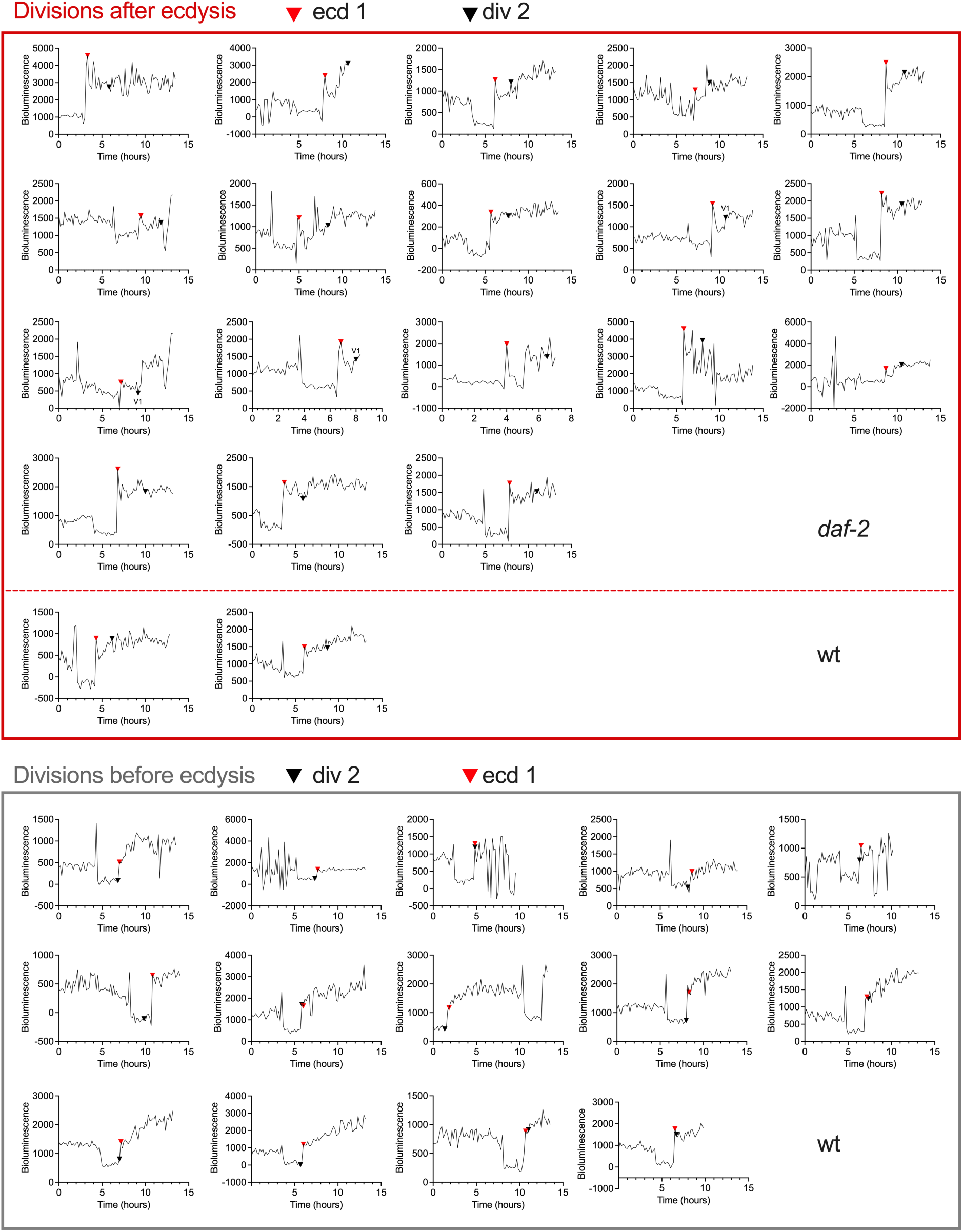
Luminescence profile from microscopy experiments for larvae categorized as seam cell divisions (div 2) ending after or before ecdysis (ecd 1). Red arrowheads signal ecdysis and black arrowheads mark V6 division, except when otherwise stated.

**Figure S5. (Relative to figure 5).**
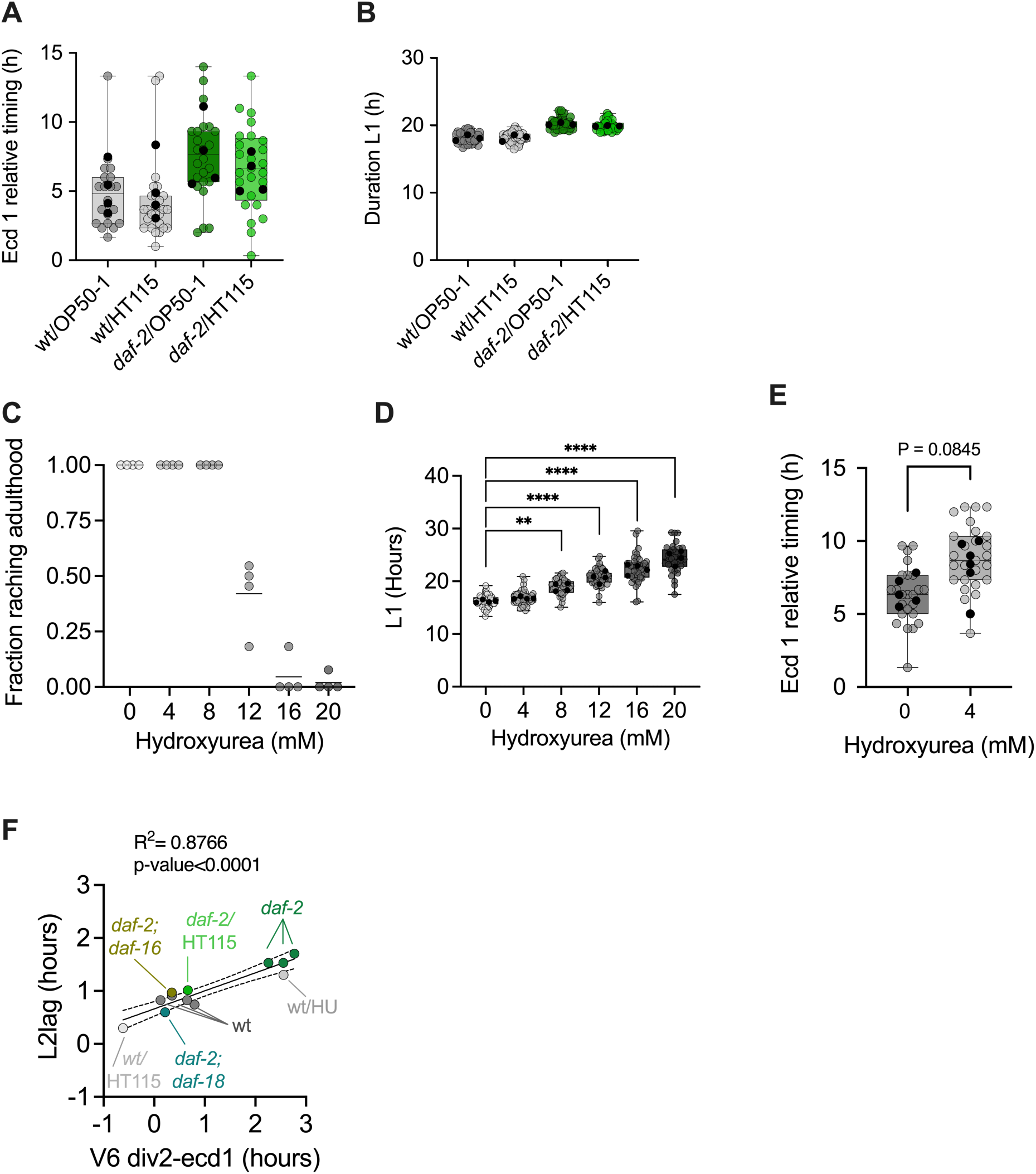
**(A)** Timing of ecdysis 1 (first release of the cuticle observed in time-lapse experiments) relative to the beginning of imaging in time-lapse experiments, for the wild type and *daf-2(e1370)* growing on OP50-1 or HT115. **(B)** Duration of the complete L1 stages, defined as the time from hatching to the end of the first molt, as inferred from the bioluminescence signal in a luminometer, for the same conditions as in A**. (C)** Fraction of animals that reach adults after treatment with different concentration of HU from L1, in four independent experiments. **(D)** Duration of the stage L1 after treatment with different concentrations of HU in four independent experiments. We performed One-way ANOVA followed by Dunnett’s multiple comparisons to test differences between each of the treatments and the untreated larvae. **(E)** Timing of ecdysis 1 (first release of the cuticle observed in time-lapse experiments) relative to the beginning of imaging in time-lapse experiments, for the wild type on 0- or 4-mM Hydroxyurea. **(F)** Correlation between L2lag and the timing of V6 seam cell division relative to ecdysis (div 2-ecd 1) for all conditions and genotypes tested. Statistics reflect the result of unpaired t-test. Differences are marked as ** p<0.005, **** p<0.0001. Grey circles show values for individual larvae and black dots show the average of each experiment. Complete statistical analysis is shown in File S1.

**S1 Table.**
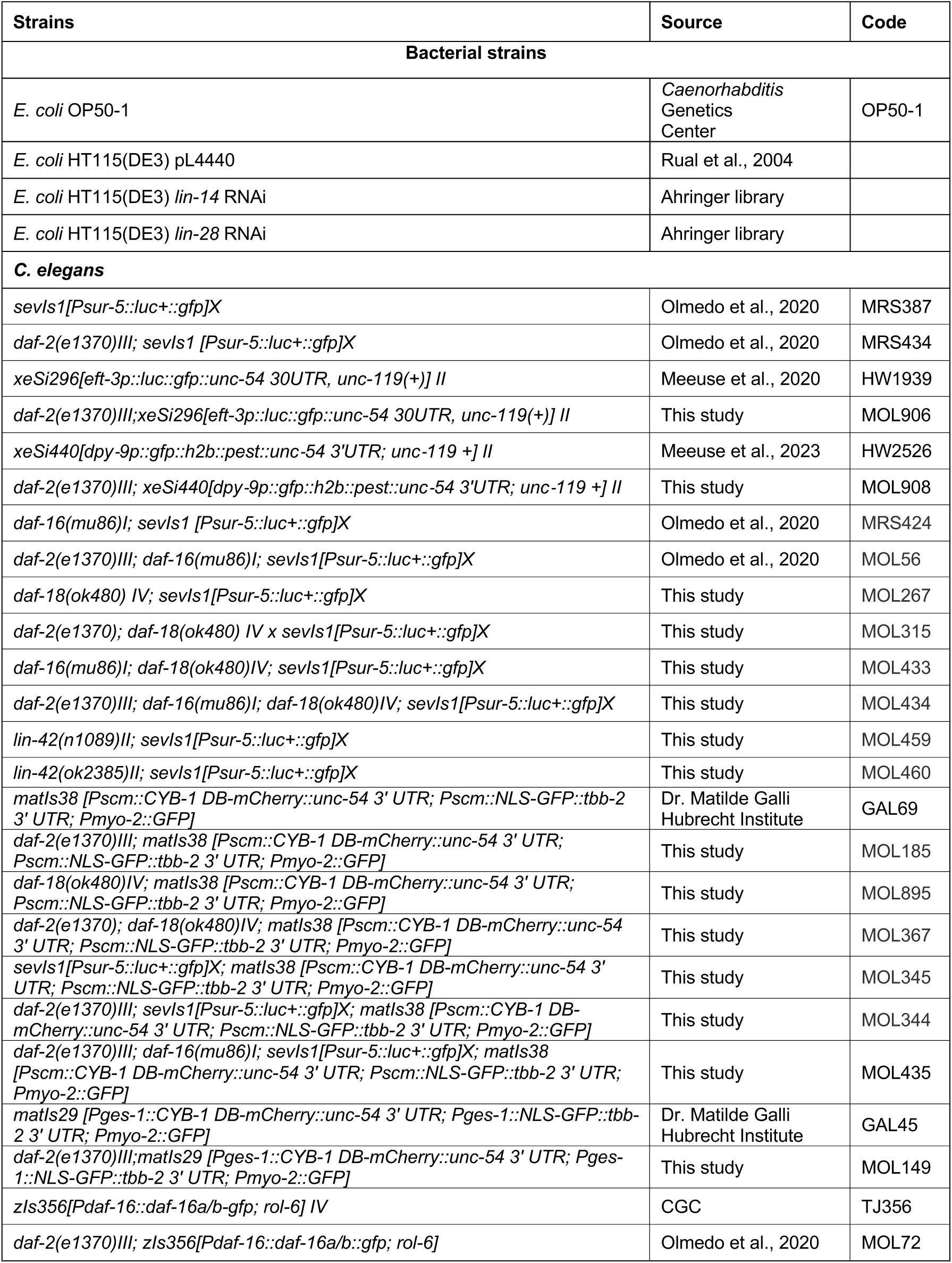
List of strains.

| Strains | Source | Code |
| --- | --- | --- |
| **Bacterial strains** |  |  |
| *E. coli* OP50-1 | *Caenorhabditis* Genetics Center | OP50-1 |
| *E. coli* HT115(DE3) pL4440 | Rual et al., 2004 |  |
| *E. coli* HT115(DE3) *lin-14* RNAi | Ahringer library |  |
| *E. coli* HT115(DE3) *lin-28* RNAi | Ahringer library |  |
| ***C. elegans*** |  |  |
| *sevls1*[*Psur-5::luc+::gfp*]X | Olmedo et al., 2020 | MRS387 |
| *daf-2(e1370)III*; *sevls1* [*Psur-5::luc+::gfp*]X | Olmedo et al., 2020 | MRS434 |
| *xeSi296*[*eft-3p::luc::gfp::unc-54 3'UTR, unc-119(+)*] II | Meeuse et al., 2020 | HW1939 |
| *daf-2(e1370)III*; *xeSi296*[*eft-3p::luc::gfp::unc-54 3'UTR, unc-119(+)*] II | This study | MOL906 |
| *xeSi440*[*dpy-9p::gfp::h2b::pest::unc-54 3'UTR; unc-119 +*] II | Meeuse et al., 2023 | HW2526 |
| *daf-2(e1370)III*; *xeSi440*[*dpy-9p::gfp::h2b::pest::unc-54 3'UTR; unc-119 +*] II | This study | MOL908 |
| *daf-16(mu86)I*; *sevls1* [*Psur-5::luc+::gfp*]X | Olmedo et al., 2020 | MRS424 |
| *daf-2(e1370)III*; *daf-16(mu86)I*; *sevls1*[*Psur-5::luc+::gfp*]X | Olmedo et al., 2020 | MOL56 |
| *daf-18(ok480) IV*; *sevls1*[*Psur-5::luc+::gfp*]X | This study | MOL267 |
| *daf-2(e1370)*; *daf-18(ok480) IV* x *sevls1*[*Psur-5::luc+::gfp*]X | This study | MOL315 |
| *daf-16(mu86)I*; *daf-18(ok480)IV*; *sevls1*[*Psur-5::luc+::gfp*]X | This study | MOL433 |
| *daf-2(e1370)III*; *daf-16(mu86)I*; *daf-18(ok480)IV*; *sevls1*[*Psur-5::luc+::gfp*]X | This study | MOL434 |
| *lin-42(n1089)II*; *sevls1*[*Psur-5::luc+::gfp*]X | This study | MOL459 |
| *lin-42(ok2385)II*; *sevls1*[*Psur-5::luc+::gfp*]X | This study | MOL460 |
| *matls38* [*Pscm::CYB-1 DB-mCherry::unc-54 3' UTR; Pscm::NLS-GFP::tbb-2 3' UTR; Pmyo-2::GFP*] | Dr. Matilde Galli Hubrecht Institute | GAL69 |
| *daf-2(e1370)III*; *matls38* [*Pscm::CYB-1 DB-mCherry::unc-54 3' UTR; Pscm::NLS-GFP::tbb-2 3' UTR; Pmyo-2::GFP*] | This study | MOL185 |
| *daf-18(ok480)IV*; *matls38* [*Pscm::CYB-1 DB-mCherry::unc-54 3' UTR; Pscm::NLS-GFP::tbb-2 3' UTR; Pmyo-2::GFP*] | This study | MOL895 |
| *daf-2(e1370)*; *daf-18(ok480)IV*; *matls38* [*Pscm::CYB-1 DB-mCherry::unc-54 3' UTR; Pscm::NLS-GFP::tbb-2 3' UTR; Pmyo-2::GFP*] | This study | MOL367 |
| *sevls1*[*Psur-5::luc+::gfp*]X; *matls38* [*Pscm::CYB-1 DB-mCherry::unc-54 3' UTR; Pscm::NLS-GFP::tbb-2 3' UTR; Pmyo-2::GFP*] | This study | MOL345 |
| *daf-2(e1370)III*; *sevls1*[*Psur-5::luc+::gfp*]X; *matls38* [*Pscm::CYB-1 DB-mCherry::unc-54 3' UTR; Pscm::NLS-GFP::tbb-2 3' UTR; Pmyo-2::GFP*] | This study | MOL344 |
| *daf-2(e1370)III*; *daf-16(mu86)I*; *sevls1*[*Psur-5::luc+::gfp*]X; *matls38* [*Pscm::CYB-1 DB-mCherry::unc-54 3' UTR; Pscm::NLS-GFP::tbb-2 3' UTR; Pmyo-2::GFP*] | This study | MOL435 |
| *matls29* [*Pges-1::CYB-1 DB-mCherry::unc-54 3' UTR; Pges-1::NLS-GFP::tbb-2 3' UTR; Pmyo-2::GFP*] | Dr. Matilde Galli Hubrecht Institute | GAL45 |
| *daf-2(e1370)III*; *matls29* [*Pges-1::CYB-1 DB-mCherry::unc-54 3' UTR; Pges-1::NLS-GFP::tbb-2 3' UTR; Pmyo-2::GFP*] | This study | MOL149 |
| *zls356*[*Pdaf-16::daf-16a/b-gfp; rol-6*] IV | CGC | TJ356 |
| *daf-2(e1370)III*; *zls356*[*Pdaf-16::daf-16a/b::gfp; rol-6*] | Olmedo et al., 2020 | MOL72 |

**S2 Table.** Strains used, number of larvae, and experimental replicates.

|  | Label | Strain | n larvae | N replicates |
| --- | --- | --- | --- | --- |
| **1E** | M1 | MRS434 | 20 | 3 |
|  | L2lag |  | 17 |  |
|  | I2 |  | 15 |  |
| **1F** | M1 | MRS434 | 17 | 5 |
|  | L2lag |  | 16 |  |
|  | I2 |  | 17 |  |
| **1G** | wt | MRS387 | 33 | 3 |
|  | *daf-2* | MRS434 | 30 |  |
| **1H** |  | MRS434 | 56 | 5 |
| **1I-J** | wt | HW2526 | 12 | 4 |
|  | *daf-2* | MOL908 | 11 |  |
| **2B,C** | wt | MRS387 | 159 | 15 |
|  | *daf-16* | MRS424 | 192 |  |
|  | *daf-2* | MRS434 | 126 |  |
|  | *daf-2;daf-16* | MOL56 | 187 |  |
| **2D,E** | wt | MRS387 | 62 | 4 |
|  | *daf-16* | MRS424 | 49 |  |
|  | *daf-18* | MOL267 | 54 |  |
|  | *daf-16;daf-18* | MOL433 | 45 |  |
|  | *daf-2* | MRS434 | 50 |  |
|  | *daf-2;daf-16* | MOL56 | 42 |  |
|  | *daf-2;daf-18* | MOL315 | 60 |  |
|  | *daf-2;daf-16;daf-18* | MOL434 | 62 |  |
| **3B** | wt/pL4440 | MRS387/ pL4440 | 33 | 3 |
|  | wt/*lin-14 RNAi* | MRS387/*lin-14 RNAi* | 34 |  |
|  | wt/*lin-28 RNAi* | MRS387/*lin-28 RNAi* | 36 |  |
|  | *daf-2*/pL4440 | MRS434/ pL4440 | 29 |  |
|  | *daf-2*/*lin-14 RNAi* | MRS434/*lin-14 RNAi* | 32 |  |
|  | *daf-2*/*lin-28 RNAi* | MRS434/*lin-28 RNAi* | 33 |  |
| **3D** | wt | MRS387 | 112 | 5 |
|  | *lin-42(n1089)* | MOL459 | 109 |  |
|  | *lin-42(ok2385)* | MOL460 | 58 |  |
| **4B** | wt | GAL69 | 15 | 4 |
|  | *daf-2* | MOL185 | 12 |  |
|  | *daf-2;daf-18* | MOL367 | 16 |  |
| **4D** | wt | GAL69 | 37 | 9 |
|  | *daf-2* | MOL185 | 26 |  |
|  | *daf-2;daf-18* | MOL367 | 34 |  |
| **4F** | wt | GAL69 | 21 | 5 |
|  | *daf-2* | MOL185 | 11 |  |
|  | *daf-2;daf-18* | MOL367 | 14 |  |
| **5A** | L2lag | MOL344 | 11 | 5 |
|  | I2 |  | 10 |  |
|  | I2 after L2lag |  | 13 |  |
| **5B** | div 2 before ecd1 | MOL345 | 14 | 9 |
|  | div 2 after ecd1 | MOL345/MOL344 | 20 (2/18) |  |
| **5C** | wt OP50-1 | GAL69 | 20 | 4 |
|  | wt HT115 |  | 26 |  |
|  | *daf-2* OP50-1 | MOL185 | 24 |  |
|  | *daf-2* HT115 |  | 24 |  |
| **5E** | wt OP50-1 | MRS387/OP50-1 | 40 | 3 |
|  | wt HT115 | MRS387/HT115 | 36 |  |
|  | *daf-2* OP50-1 | MRS434/OP50-1 | 41 |  |
|  | *daf-2* HT115 | MRS434/HT115 | 38 |  |
| **5F** | 0 mM HU | GAL69 | 23 | 6 |
|  | 4 mM HU |  | 21 |  |
| **5H** | 0 mM HU | MRS387 | 44 | 4 |
|  | 4 mM HU |  | 43 |  |
| **6A** | 12 °C | MRS387 | 38 | 3 |
|  | 16 °C |  | 61 | 3 |
|  | 20 °C |  | 40 | 3 |
|  | 22 °C |  | 38 | 3 |
| **6B,C** | 20 °C | GAL69 | 15 | 4 |
|  | 22.5 °C |  | 15 | 4 |
| **6D** | 12 °C | MRS434 | 46 | 3 |
|  | 16 °C |  | 61 | 3 |
|  | 20 °C |  | 39 | 3 |
|  | 22 °C |  | 38 | 3 |
| **6E,F** | 20 °C | MOL185 | 13 | 4 |
|  | 22.5 °C |  | 18 | 4 |

